# Bayesian inference of the gene expression states of single cells from scRNA-seq data

**DOI:** 10.1101/2019.12.28.889956

**Authors:** Jérémie Breda, Mihaela Zavolan, Erik van Nimwegen

## Abstract

In spite of a large investment in the development of methodologies for analysis of single-cell RNA-seq data, there is still little agreement on how to best normalize such data, i.e. how to quantify gene expression states of single cells from such data. Starting from a few basic requirements such as that inferred expression states should correct for both intrinsic biological fluctuations and measurement noise, and that changes in expression state should be measured in terms of fold-changes rather than changes in absolute levels, we here derive a unique Bayesian procedure for normalizing single-cell RNA-seq data from first principles. Our implementation of this normalization procedure, called Sanity (SAmpling Noise corrected Inference of Transcription activitY), estimates log expression values and associated errors bars directly from raw UMI counts without any tunable parameters.

Comparison of Sanity with other recent normalization methods on a selection of scRNA-seq datasets shows that Sanity outperforms other methods on basic downstream processing tasks such as clustering cells into subtypes and identification of differentially expressed genes. More importantly, we show that all other normalization methods present severely distorted pictures of the data. By failing to account for biological and technical Poisson noise, many methods systematically predict the lowest expressed genes to be most variable in expression, whereas in reality these genes provide least evidence of true biological variability. In addition, by confounding noise removal with lower-dimensional representation of the data, many methods introduce strong spurious correlations of expression levels with the total UMI count of each cell as well as spurious co-expression of genes.

## Introduction

In the past decade much effort has been invested in adapting methods for quantifying transcriptome and epigenome state on a genome-wide scale to the single-cell level. This has led to a large number of new methods that are starting to make it possible to track the states of single cells across tissues and embryos as they are developing, measuring transcriptomes, chromatin state, chromatin conformation, and cell lineages, sometimes in parallel [1–22]. Many in the field believe that these single-cell methods will revolutionize our understanding of the ways in which cell fate, cell identity and developmental processes are regulated, and major consortia are starting to form that aim to comprehensively map single-cells in model organisms [23, 24].

In order to fullfil the promise of these single-cell measurement technologies, it will be crucial that computational methods are available that unambiguously extract what the raw measurements say about the state of the single cells in terms of concrete physical quantities. We not only want to be able to integrate results of single-cell RNA-seq (scRNA-seq) measurements, which we will focus on in this work, from different labs using different protocols, but also across measurements from entirely different measurement technologies such as FISH (e.g. [25]). In order to make that possible, the expression values that we extract from scRNA-seq data should correspond to physically meaningful quantities that can be directly compared with measurements of the same quantities made with other experimental methods. In addition, the estimated values of the concrete physical quantities should follow directly from the experimental data together with as small a number of additional assumptions as possible, and not depend on arbitrary parameters that the user can set at will. Moreover, in order to be able to determine when different measurements are mutually consistent, all estimates should be accompanied by meaningful error bars.

However, although there has been a veritable explosion of scRNA-seq analysis tools in recent years, there has been almost no attention given to satisfying these objectives. Instead of there being a small number of transparent methods that provide unambiguous estimates of quantities with clear physical interpretation, we find a large number of *ad hoc* methods that apply highly complex transformations to the data to perform combinations of tasks including imputation/normalization, clustering, dimensionality reduction, pseudo-time and trajectory inference, and visualization. These methods typically have many tunable parameters, produce outputs in highly abstract spaces that lack clear biological meaning, and are often even stochastic, such that different runs on the same data with the same parameters result in different output. For example, probably the most popular tools for visualizing scRNA-seq data are t-SNE [26] and UMAP [27], which are both stochastic, involve several parameters, and position cells in a lower dimensional space whose dimensions lack biological interpretation.

We here focus on the relatively basic task of normalization/imputation of single-cell gene expression states from raw scRNA-seq transcript counts. Using only minimal assumptions we derive from first principles a Bayesian method that corrects not only for the finite sampling associated with the capture and sequencing of mRNAs, but also for the Poisson noise inherent in the gene expression process itself. Our method, which we call Sanity (SAmpling Noise corrected Inference of Transcription activitY) is deterministic, has zero tunable parameters, and provides error-bars for all its estimates.

After motivating and explaining our method, we compare Sanity with a selection of popular methods for imputation/normalization from the recent literature and show that only Sanity can meaningfully remove Poisson sampling fluctuations and infer the true variation in gene expression intensity of each gene across cells. In addition, we show that all other methods we tested introduce severe distortions of the data such as inducing strong correlations between expression estimates and total UMI count of cells, or inferring strong co-expression between large numbers of genes when none is evident in the data. In addition, we show that Sanity’s estimated expression levels outcompete those of other methods on both downstream clustering and differential expression tasks.

## Methods

### A Bayesian method for inferring gene expression states from count data

We first motivate and explain how we represent gene expression states of single cells, and what concrete physical quantities these gene expression states correspond to. After that, we introduce our method’s probabilistic model of a scRNA-seq experiment, calculating how the expression state of the cell determines the probabilities of obtaining particular raw transcript counts, and then discuss how we solve the Bayesian model and the outputs that the method provides.

### Defining gene expression states

For any given cell *c*, we want to represent its ‘gene expression state’ by a vector 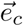, whose components *e*_*gc*_ quantify how strongly each gene *g* is expressed in the cell. We want these gene expression states to satisfy two basic criteria. First, these gene expression states should have concrete physical interpretation. Second, for downstream processing we want that the differences *e*_*gc*_ − *e*_*gc*′_ meaningfully reflect the change in expression of gene *g* between cells *c* and *c*′ such that the Euclidean distance 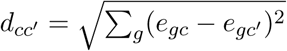 between two cells *c* and *c* meaningfully reflects the difference in their gene expression states.

One might think that we could simply take the vector 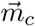 of the actual number of mRNAs *m*_*gc*_ that exist in cell *c* for each gene *g* as the gene epression state of the cell. However, even for cells in the same gene expression state, the number of mRNAs will exhibit stochastic fluctuations. Imagine a gene that is transcribed at a constant rate *λ* in every cell, and with a constant rate of mRNA decay *μ* in every cell. The actual number of mRNAs *m* across cells will then follow a Poisson distribution with mean *a* = *λ/μ* which we call its ‘transcription activity’. That is, the probability to find *m* mRNAs is *P*_*m*_ = *a*^*m*^*e*^−*a*^*/m*! which has mean ⟨*m*⟩ = *a* and variance var(*m*) = *a*. Thus, instead of assuming any change in mRNA number *m* reflects a change in gene expression state, it makes more sense to identify changes in gene expression state with changes in the transcription activity *a*.

Note that mRNA numbers will show Poisson fluctuations in much more general situations than constant rates of transcription and decay [28]. Imagine that, in a particular cell *c*, both the rate of transcription and mRNA decay of a given gene *g* has fluctuated in some arbitrary way in time, with *λ*_*gc*_(*t*) the transcription rate a time *t* in the past, and *μ*_*gc*_(*t*) the mRNA decay rate a time *t* in the past. The expected number of mRNAs ⟨*m*_*gc*_⟩ is then given by the transcription activity

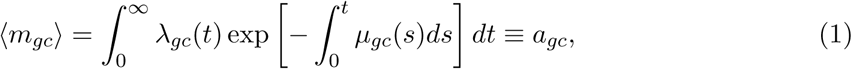

which is a weighted average of the transcription rate of the gene in the recent past, i.e. on the time-scale that its mRNAs have turned over. Given this expected mRNA number *a*_*gc*_ the distribution of the actual number of mRNAs *m*_*gc*_ is still Poisson. That is the probability to obtain *m*_*gc*_ mRNAs is

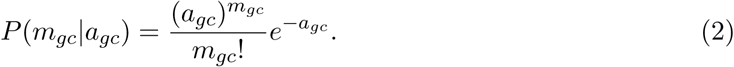

We thus propose that we should use changes in vectors of transcription activity 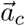 to represent changes in gene expression state.

In addition, we propose to characterize the gene expression state of a cell not by the vector 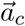 of absolute transcription activities *a*_*gc*_, but by the vector 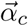 of *relative* transcription activities, with

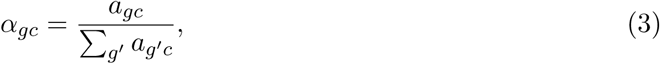

which we will refer to as *transcription quotients*. First, it has been shown that, as cell volume increases, cells globally upregulate transcription to maintain approximately constant mRNA concentration [29] so that transcriptional activities *a*_*gc*_ of all genes are generally expected to scale with cell volume. We argue that a global change in transcriptional activities by a common factor *c*, i.e. *a*_*gc*_ → *ca*_*gc*_ for all genes, does not correspond to a change in gene expression state, but just to a change in cell size. Second, it is well known that, in current scRNA-seq protocols, the rate of capture and sequencing of mRNAs varies significantly across cells [30, 31] so that there is only a weak quantitative relationship between the total number of sequenced mRNA molecules and the true total mRNA content of cells. Although it is possible to estimate capture and sequencing efficiencies, at least to some extent, using RNA spike-in controls [30, 32], most experiments are performed without such controls. Therefore, for most scRNA-seq datasets it is unclear to what extent variations in total sequenced mRNAs across cells represent biological variability, as opposed to technical variability. Consequently, transcription quotients can generally be much more accurately estimated than absolute transcription activities, because they do not directly depend on capture efficiency. Note that quantifying gene expression by quotients, i.e. transcripts per million transcripts, is also the standard approach in bulk RNA-seq experiments.

Finally, we note that if we were to use differences in transcription quotients of mRNAs *α*_*gc*_ − *α*_*gc*′_ to quantify the change in expression of gene *g* between cells *c* and *c*′, then this change would be proportional to overall expression level of the gene. That is, a change from 20 to 40 transcripts per million would be considered ten times as large as a change from 2 to 4 transcripts per million. Since the early days of transcriptomics it has been observed [33] that, as would be expected from the multiplicative effects of fluctuations in rates of various biochemical reactions [34], the relative expression levels of genes in a sample follows a roughly log-normal distribution that covers several orders of magnitude. Consequently, if we were to quantify expression changes directly by the changes *α*_*gc*_ − *α*_*gc*′_, the expression changes between two cells would be completely dominated by those of the highest expressed genes. Therefore, it has long become standard to instead use *logarithms* of the expression levels. Thus, we propose to quantify the gene expression state of a cell by the *logarithms of the transcription quotients* (LTQs) log(*α*_*gc*_) so that a *x*-fold change in quotient *α*_*gc*_ → *α*_*gc*′_ = *xα*_*gc*_ corresponds to the same additive change log(*α*_*gc*_) → log(*α*_*gc*_) + log(*x*) in LTQ, independent of the absolute value of the quotient *α*_*gc*_. In summary, we propose to characterize the gene expression state of a cell *c* by a vector of LTQs log(*α*_*gc*_).

### A probabilistic model for a scRNA-seq experiment

The initial steps of scRNA-seq analysis involve basic processing of the raw sequencing reads such as quality control of the reads, identification of the various barcodes that identify the library, the individual cell, the unique mRNA molecule (if available), and mapping each read to the corresponding genome or transcriptome. The methods used in these steps are similar to methods used for bulk RNA-seq and ChIP-seq and have matured to the point that there are accepted methods and little variability in the results from commonly used tools, e.g. [35–38].

The introduction of unique molecule identifiers (UMIs) [39] was an important development in scRNA-seq technology in that it avoids noise in expression measurements due to fluctuations in PCR amplification, and determineds the number of unique mRNA molecules that were captured for each mRNA. Since only protocols that incorporate UMIs allow for a realistic modeling of the statistics of the measurement noise, we will here focus on scRNA-seq protocols that use UMIs.

After the basic processing of the raw data has been performed, the data will consist of a matrix of integers *n*_*gc*_ giving the number of captured mRNA molecules for each gene *g* in each cell *c*. The key assumption of our probabilistic model is that, in a scRNA-seq experiment, each mRNA molecule in a given cell *c* has a probability *p*_*c*_ to be captured and sequenced. This capture probability *p*_*c*_, which varies from cell to cell, has been estimated to be in the range of 10 to 15% [40] and up to 30% with the most recent protocols [41]. Under this assumption, the probability of the observed UMI counts *n*_*gc*_ in cell *c* given the transcription quotients *α*_*gc*_ is still given by a product of Poisson distributions (see Supplementary Methods)

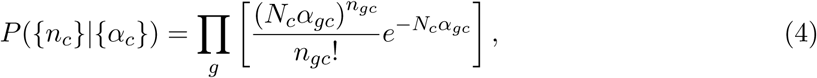

where {*n*_*c*_} is the set of UMI counts in cell *c*, {*α*_*c*_} the set of transcription quotients in cell *c*, and *N*_*c*_ the total number of UMIs in cell *c*. Crucially, we see that the convolution of the biological Poisson noise and the sampling noise introduced by the scRNA-seq measurement together still lead to a simple Poisson distribution in terms of the transcription quotients *α*_*gc*_.

### Prior probabilities and the Bayesian solution

In order to estimate the log(*α*_*gc*_) from the observed UMI counts *n*_*gc*_ a final ingredient that we need is to define a prior distribution over these LTQs. As we aim to minimize the number of assumptions that our inference makes, our model will not assume any dependence structure between the LTQs of different genes, i.e. we will not assume that the gene expression data derives from a low-dimensional manifold. We will also not assume that the LTQs follow a particular distribution. The only thing we will assume is that, for each gene, the prior distribution of LTQs log(*α*_*gc*_) can be characterized by its mean *μ*_*g*_ and variance *v*_*g*_. We rewrite the transcription quotients *α*_*gc*_ in terms of an average quotient *α*_*g*_ and a cell-specific log fold-change *δ*_*gc*_, i.e. 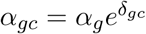. With that reparametrization, the mean *μ*_*g*_ equals log(*α*_*g*_) and the *δ*_*gc*_ derive from a prior probability distribution with mean zero and variance *v*_*g*_. Given that we only specify the variance of the distribution of the *δ*_*gc*_ to be *v*_*g*_, we choose the maximum entropy distribution [42] consistent with this constraint, which is a Gaussian distribution. Importantly, this does not mean that we assume that the log fold-changes *δ*_*gc*_ follow a Gaussian distribution. It just assumes that, before seeing any of the data, we assume the *δ*_*gc*_ are taken from the broadest, least assuming distribution consistent with some (unknown) variance *v*_*g*_.

In the Supplementary Methods we derive in detail how this model can be solved to estimate, for each gene *g*:

1. The mean LTQ *μ*_*g*_ and its error-bar *δμ*_*g*_.
2. The estimated variance *v*_*g*_ of the changes in LTQs *δ*_*gc*_ across cells.
3. For each cell *c*, the estimated LTQ 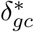 and an error-bar *ϵ*_*gc*_ on this LTQ.

Note that the LTQs 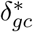 provide estimates for how much the transcription and decay rates of each gene *g* in cell *c* differ from their average rates, and thus correct for both the intrinsic biological Poisson fluctuations as well as the finite sampling fluctuations inherent in the scRNA-seq measurement.

### Alternative methods for scRNA-seq normalization

To assess the performance of Sanity we will compare it with a number of other methods for normalization/imputation from scRNA-seq data. Apart from a number of other tools from the recent literature, we also include two basic normalization procedures that are widely used. First, the simplest approach to estimating gene expression levels *e*_*gc*_ from scRNA-seq data is to simply log-transform the observed number of UMIs *n*_*gc*_ after adding a *pseudocount p* in order to avoid problems with zero counts *n*_*gc*_ = 0, i.e.

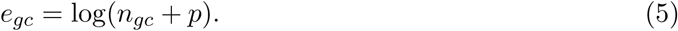

A typical choice for the pseudo-count is *p* = 1, because it attenuates fluctuations in *n*_*gc*_ on the order of magnitude corresponding to the resolution of the experimental measurements. We will refer to this normalization, with *p* = 1, as the *RawCounts* normalization, since it essentially just log-transforms the raw counts.

However, the total number *N*_*c*_ of mRNAs captured and sequenced from an individual cell *c* can vary substantially due to fluctuations in capture efficiency and sequencing depth, as well as changes in cell size. Consequently, the RawCounts procedure introduces artificial correlations between the expression levels *e*_*gc*_ and the total number of UMIs *N*_*c*_ that were sequenced in the cell *c*. Thus, the most commonly used normalization approach is to first divide the rawcounts *n*_*gc*_ by the total counts *N*_*c*_ and then multiply by a typical total count *N* before adding a pseudocount and log transforming, i.e.:

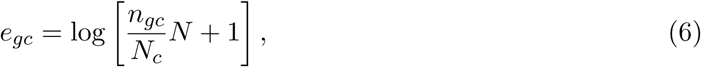

where we will take for the typical total count *N* the median of the counts *N*_*c*_ across all cells. In a slight abuse of terminology, we will call this normalization the *TPM* normalization because of its close connection to the transcripts per million normalization used in bulk RNA-seq (which corresponds to setting *N* = 10^6^).

Beyond these two simple normalization methods, we compare Sanity’s performance with that of the following recently published tools:

1. DCA [43], which uses a deep learning based autoenconder.
2. MAGIC [44], which uses diffusion of measured gene expression states between cells with similar expression profiles.
3. SAVER [45], which assumes negative binomial counts distributions *n*_*gc*_ and models the underlying rates using Poisson LASSO regression with the expression levels of other genes.
4. scImpute [46], which focuses mainly on correcting ‘dropouts’, i.e. datapoints for which *n*_*gc*_ = 0.
5. scVI [47], which uses a deep neural network based autoencoder.

Note that, with the exception of scImpute, all these methods seek to normalize the expression levels for the total UMI count per cell, and seek to remove noise by using lower dimensional representations of the input counts *n*_*gc*_.

We used default parameters for all these methods and, since all methods report expression values in linear space, we log-transformed all expression values. MAGIC sometimes reports 0 or negative values and, as suggested by the authors, we first set all negative values to 0 and then add a pseudocount of 1 to all expression values (including the nonzero ones) before log-transforming. Similarly, scImpute reports some zero values and we added a pseudocount of 1 to all these.

### Test datasets

To comprehensively assess the performance of the different methods we used a collection of datasets for which annotation of the sequenced cell types was available. The datasets we used were (labelled by the first author of the publication):

1. *Grün*: 160 mouse embryonic stem cells and 160 corresponding aliquots consisting of, 80 cells from culture in 2i medium, 80 cells from culture in serum, and 80 aliquots for each condition that were created by pooling cells together, and then splitting the pool into single-cell mRNA equivalents [30].
2. *Zeisel*: 3’005 cells from the somatosensory cortex and from the CA1 region of the mouse hippocampus, annotated into 7 cell types [48].
3. *Baron*: 1’937 human pancreatic cells annotated into 14 cell types [49].
4. *Chen*: 14,437 adult mouse hypothalamus cells annotated into 15 clusters [50].
5. Three datasets from *LaManno* [51]:
  a. *LaManno/Embryo*: 1’977 ventral mid-brain cells from human embryo annotated into 25 classes.
  b. *LaManno/ES*: 1’715 human embryonic stem cells annotated into 17 classes.
  c. *LaManno/MouseEmbryo*: 1’907 ventral mid-brain cells from mouse embryo annotated into 26 classes.

In addition to these real datasets we also constructed one simulated dataset as described in the Supplementary Methods. The parameters of the simulation were chosen so as to mimic the statistics of the *Baron* dataset (see Fig. S11).

## Results

### Sanity accurately corrects for Poisson fluctuations to identify true variance in gene expression

A key aim of Sanity’s normalization is to correct for both biological and technical sampling fluctuations in order to quantify the true biological variation in expression of each gene across cells. Testing this is challenging because the true expression variability of each gene is generally unknown. To address this we used a simulated dataset for which the true expression variability of each gene is known, on the one hand, and analyzed a carefully designed study of mouse embryonic stem cells (ESCs) from *Grün et al* [30] on the other hand. In this study, mouse ESCs were culured in both 2i and serum conditions and apart from scRNA-seq measurements on these cells, the same measurement protocol was applied to single-cell equivalent *aliquots* from pooled RNA of multiple cells. Since all aliquots were sampled from the same pool, there is no biological expression variation in this dataset at all, and the expression variation in these aliquots derives solely from technical sampling noise. In addition, the ESCs are highly homogeneous so that also little true expression variation is expected for ESCs in the same condition, and the main expression differences are expected between cells in the 2 different culture conditions.

The amount of expression variability of a gene across a set of cells can be quantified by its coefficient of variation CV, i.e. the ratio of the standard-deviation and the mean of its expression levels. Figure 1A shows box-whisker plots of the distribution of CVs, for each of the 4 datasets, as calculated from the (non log-transformed) expression estimates of each of the normalization methods. Ideally the methods should infer that there is no true variability at all for the aliquots, and relatively little variability for the ESCs. In addition, it is known that variability in 2i conditions is smaller than in serum [30], so that we expect larger CVs in serum. Although, with the exception of scVI, all methods infer that the CVs are larger in serum than in 2i, and smaller for the aliquots than for the cells, the CVs that Sanity infers are at least twofold lower than those of all other methods, and only Sanity infers that the CV is less than 10% for the large majority of the aliquots. Of the other methods, MAGIC and SAVER show distributions of CVs that, while generally larger, are closest to those inferred by Sanity. All other methods show distributions of CVs that are at odds with our prior information in one way or another. For example, due to the Poisson noise, the simple RawCounts, TPM, and scImpute methods infer CVs of at least 0.5 for the large majority of genes in both cells and aliquots. DCA infers very similar distributions of CVs for the ESCs and aliquots and, finally, scVI shows unrealistically high CVs for all genes in both ESCs and aliquots.

**Figure 1:**
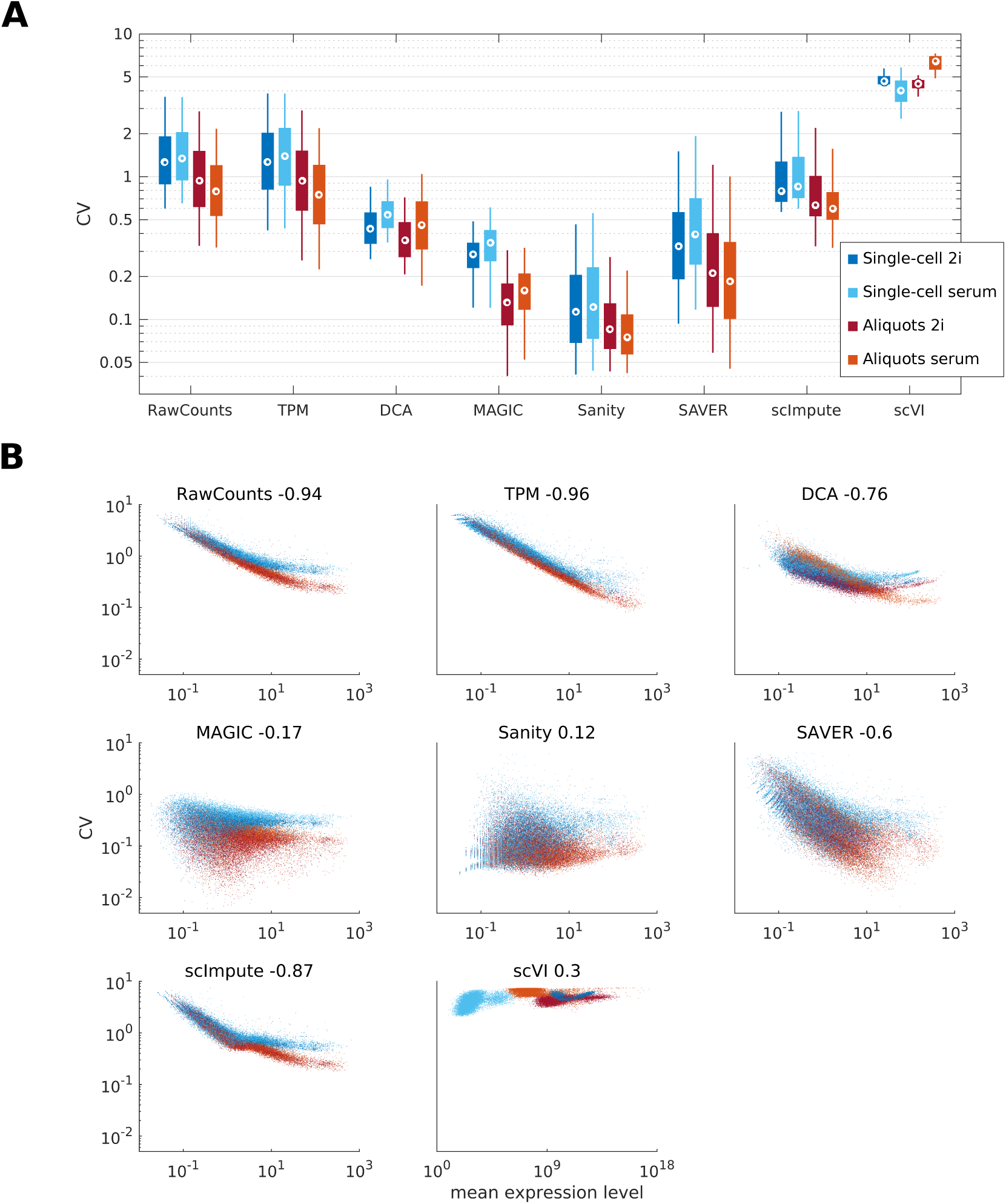
**A**: Box-whisker plots showing the medium (circle) as well as the 5th, 25th, 75th, and 95th quantiles of the distribution of gene expression levels for each of the 4 datasets (see legend) as inferred by each of the normalization methods. **B** Scatter plots of CV (standard-deviation divided by mean) for all genes in each of the 4 datasets (colors as in panel A) as inferred by each of the normalization methods. The Pearson correlation coefficient between log *CV* and log mean is shown on top of each plot. The axis are shown on a logarithmic scale and are kept similar across panels, except for *scVI* where the mean expression values are on a very different scale from those of the other methods.

The ability for methods to correct for sampling noise can be assessed most clearly by plotting the CV of each gene as a function of its mean expression (Fig. 1B). As is well appreciated in the scRNA-seq literature, e.g. [32], because the variance of a Poisson distribution is equal to its mean, Poisson sampling fluctuations add a term 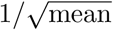 to the CV. Because most genes have low absolute expression values, the CV is dominated by this term for most genes, leading to a strong negative correlation between mean expression and CV. Indeed, the simple RawCounts and TPM methods show an almost perfect negative correlation between CV and mean, showing that sampling noise dominates the observed variability for all but the highest expressed genes. Ideally, the normalization would correct for the Poisson contribution to the CV, and in principle we would not expect to see a systematic correlation between mean expression and the normalized CV. Indeed, for Sanity the normalized data does not exhibit any correlation between CV and mean. However, with the exception of MAGIC and scVI, a strong negative correlation between CV and mean remains for all other methods, showing that even after normalization the expression variability is dominated by Poisson noise for many genes. We also note that scVI exhibits rather unnatural distributions of CV and mean, with consistently high CV and strongly varying means across datasets. These observations do not only apply to the dataset of [30], but are observed for all test datasets we considered (Suppl. Fig. S1).

We next constructed a simulated dataset (see Supplementary Methods) in which the total number of UMI per cell and mean expression levels where chosen to match those of the dataset of Baron et al. [49]. Each gene was assigned a random true variance in log gene expression, its true expression values were sampled from a Gaussian distribution with corresponding variance, and finally Poisson noise was added to these true expression values. Since, for this simulated dataset, we know exactly the true variability in gene expression for each gene, we directly compared the inferred CV with the true CV used in the simulation (Fig. 2). For the simple TPM and RawCounts methods, there is actually a good correlation between true CV and inferred CV for very highly expressed genes, confirming that for the highest expressed genes the observed CV in the data matches the true CV. However, for the large majority of genes, the Poisson noise causes the inferred CV to be much higher than the true CV, so that there is ultimately almost no correlation between true and observed CVs across the entire set of genes. For most of the other methods, there is very little relation between the true and inferred CVs. MAGIC and DCA predict much lower CVs than the true CVs, scVI predicts consistently high CVs, and there is no correlation between the true and predicted CVs for any of these methods. Only Sanity and SAVER show a good match between the true and inferred CVs across most of the genes. Notably, Sanity accurately estimates CVs for all highly expressed or highly variable genes. For low expressed genes, where there is not sufficient data to reliably detect the true expression variability of a gene, Sanity conservatively infers that the true expression variability is low and these genes will therefore not significantly contribute to any downstream analysis of expression variability across cells. Although SAVER’s inferred CVs are reasonable for most genes, they are clearly less accurate than Sanity’s predictions, and for a subset of low expression genes SAVER strongly overestimates the CV.

**Figure 2:**
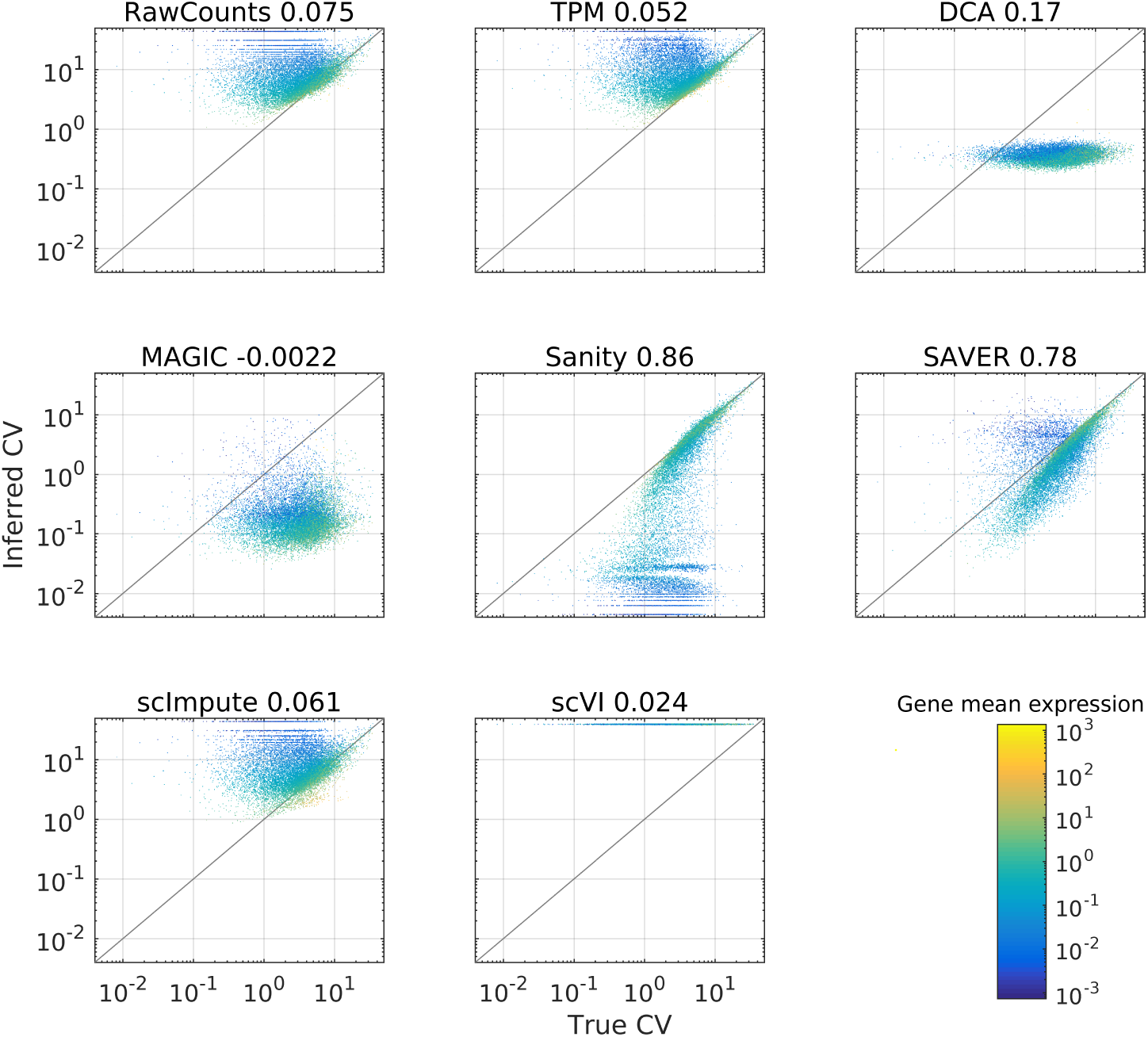
Comparison of the true CVs and those inferred by each of the normalization methods on the simulated dataset. Each panel shows a scatter plot of the true CV (horizontal axis) the CV as inferred by the normalization method (vertical axis) across genes. The color of each datapoint shows the mean expression level of the gene (total UMI in the dataset, see colorbar). The Pearson correlation between the inferred CVs and the true CVs is shown on top of each panel.

In summary, these results show that Sanity is the only normalization method that can reliably correct for the Poisson sampling noise to quantify the true expression variability of each gene.

### Many normalization methods introduce spurious correlations with library size

Due to variations in cell size, mRNA capture efficiency, and sequencing depth, the total number of captured UMIs can fluctuate significantly from cell to cell. Therefore, most scRNA-seq processing methods involve normalize the expression levels of genes in a given cell for the total number of mRNAs (i.e. UMIs) that were sequenced for that cell. For example, whereas the simple RawCounts procedure does *not* correct for total UMI counts per cell, the simple TPM procedure normalizes for total UMI count by dividing the observed counts for each gene by this total count. With the exception of scImpute, all other methods also include methods to normalize for total UMI count.

To investigate the effects of the normalization for total UMI count we calculated, for each method and each gene, the Pearson correlation between the inferred log expression levels and the logarithm of the total UMI count across cells. Using the Zeisel dataset as an example, Fig. 3 shows the distribution of Pearson correlations for each of the methods as well as the raw scatters of the normalized expression levels as a function of log total UMI count for one example gene (*Zbed3*). Starting with the simple RawCounts method we see, as expected from the fact that this method does not normalize for total UMI count, that for most genes there is a positive correlation between total UMI count of a cell and the expression level of the gene in that cell. The scImpute method shows similar correlations with total UMI count which is consistent with the fact that this method does not normalize for total UMI count either. In contrast both the simple TPM method, and especially Sanity, remove this correlation, confirming that these methods successfully normalize for the fluctuations in total UMI count across cells.

**Figure 3:**
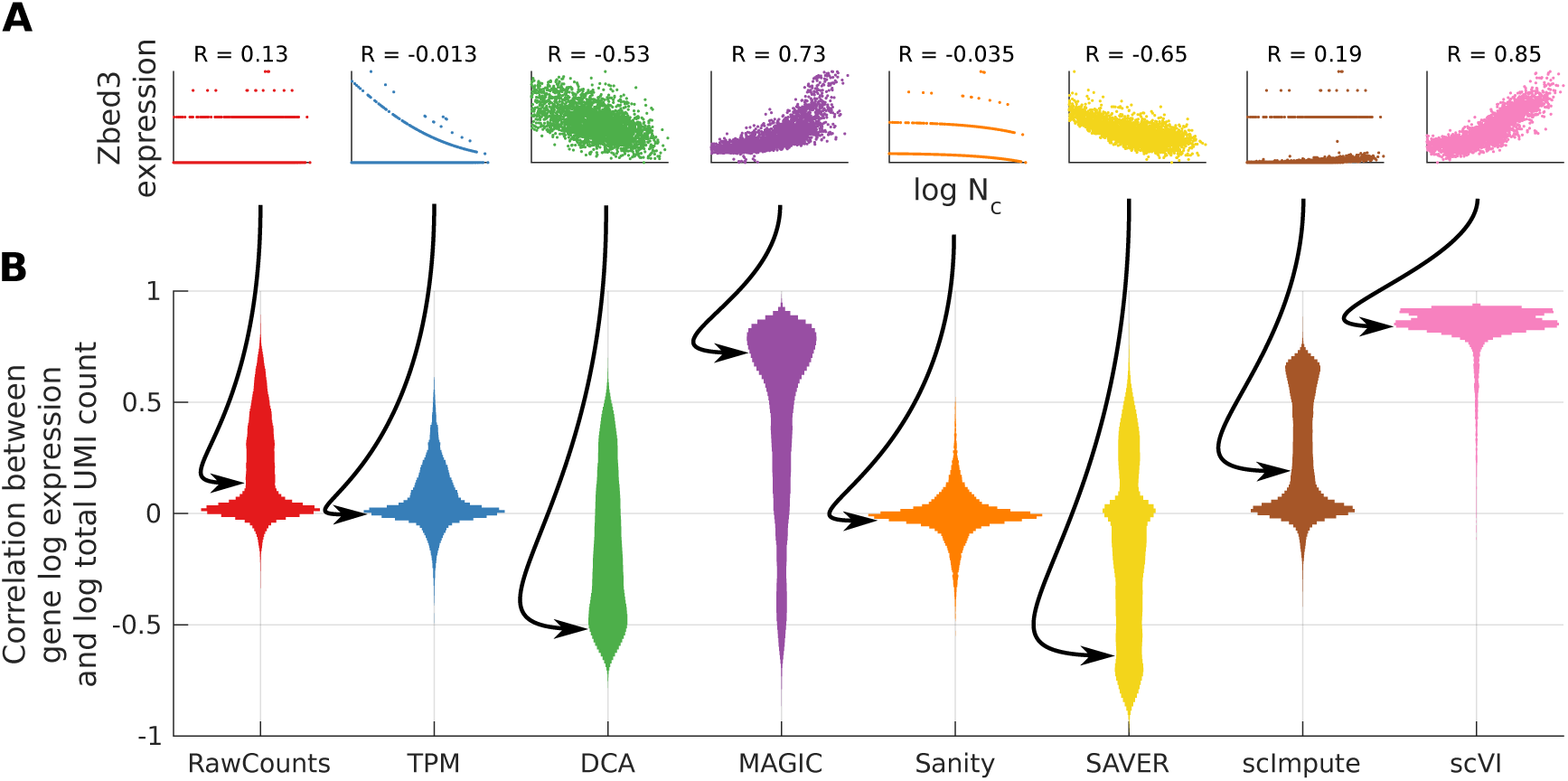
**A**: Scatter plots of the normalized log expression level of the example gene (*Zbed3*) versus the logarithm of the total UMI count log(*N*_*c*_) across cells for each of the methods. The Pearson correlation of the dependence is shown above each panel. **B**: Violin plots of the distribution of correlation coefficients between the inferred log expression levels of genes and the log of total UMI count per cell, for the *Zeisel* dataset. Different colors correspond to the different methods, which are indicated below.

We were very surprised to see that for all other methods, rather than removing correlations with total UMI count, the normalized expression levels show even stronger correlations with total UMI count. DCA, SAVER, and MAGIC show a very wide distribution of correlation coefficients with predominantly negative correlations for DCA and SAVER, and predominantly positive correlations for MAGIC. The situation is even more dramatic for scVI which infers that the expression levels of essentially *all* genes are highly correlated with total UMI count. The scatters with the predicted gene expression levels for the gene *Zbed3* as a function of log total UMI count log(*N*_*c*_) illustrate how dramatically the various normalization methods transform the input data. The RawCounts show that this gene is fairly low expressed, with either 0 or 1 UMI observed in most cells, and that there is a slightly higher chance to observe one or two UMIs when the total UMI count *N*_*c*_ is larger. However, DCA, MAGIC, SAVER, and scVI completely transform this input data into a scatter of continuously varying expression levels that either correlate strongly negatively (DCA, SAVER) or strongly positively (MAGIC, scVI) with total UMI count. These observations again generalize to all other datasets as shown in Suppl. Fig. S2. In conclusion, only TPM and Sanity reliably normalize for variations in total UMI count, and most other methods introduce strong spurious correlations with total UMI count.

### Most normalization methods spuriously infer co-expression between many pairs of genes

One of the most common downstream analyses that are applied to transcriptome data is the identification of co-expressed genes, for example for identifying co-regulated pathways or regulatory modules. In order to perform such co-expression analysis, it is crucial that the pairwise correlations of the normalized expression profiles across genes accurately reflect the co-expression evidence in the data. In order to compare co-expression information across methods we calculated, for each method and for every pair of genes, the Pearson correlation of their normalized expression levels. We then compared these pairwise correlation coefficients across the various methods.

Using the Baron dataset as an example, Fig. 4A shows a scatter of the pairwise correlations as inferred by Sanity and the simple TPM method for all pairs of genes. We see that the inferred pairwise correlations by-and-large agree between the two methods, i.e. most points fall along the diagonal, and there are almost no pairs where the two methods strongly disagree on the strength of the correlation. As shown in Suppl. Fig. S3, this also holds for the comparison of Sanity’s pairwise correlations with those of RawCounts and scImpute. However, a very different pattern is observed for the comparison of Sanity with MAGIC (Fig. 4C). For many of the pairs of genes for which Sanity infers no co-expression, i.e. zero correlation, MAGIC infers a broad range of correlations running from almost perfect anti-correlation, to perfect correlation. To assess whether the raw data are more consistent with Sanity’s or MAGIC’s pairwise correlations, we first focused on a subset of 4360 pairs of genes within the red rectangle of Fig. 4C, for which MAGIC predicts nearly perfect correlation (*r* > 0.975) whereas Sanity predicted none (−0.03 < *r* < 0.005). Summing across all 4360 pairs and all cells, we counted the total number of times *n*_*i,j*_ for which *i* UMIs were observed for the first gene and *j* for the second. Strikingly, there was not a single example for which both *i* and *j* are larger than zero (Fig. 4D). That is, although MAGIC infers that these 4360 pairs of genes are almost perfectly co-expressed, *none* of them are ever observed to be present at the same time in *any* cell. In contrast, for the small set of pairs for which Sanity infers significant co-expression whereas MAGIC does not (magenta box in Fig. 4C), we do generally find evidence of co-expression (Fig. 4E). As shown in Suppl. Fig. S3, the same pattern is observed for the comparisons of Sanity’s pairwise correlations with those of DCA and SAVER. That is, MAGIC, DCA, and SAVER all infer large numbers of highly correlated or anti-correlated pairs of genes, whereas there is no evidence at all in the raw data that these pairs of genes are co-expressed. The pairwise correlations predicted by scVI show even more pathological behavior, i.e. scVI predicts that *all* pairs of genes are significantly co-expressed (Fig. 4B).

**Figure 4:**
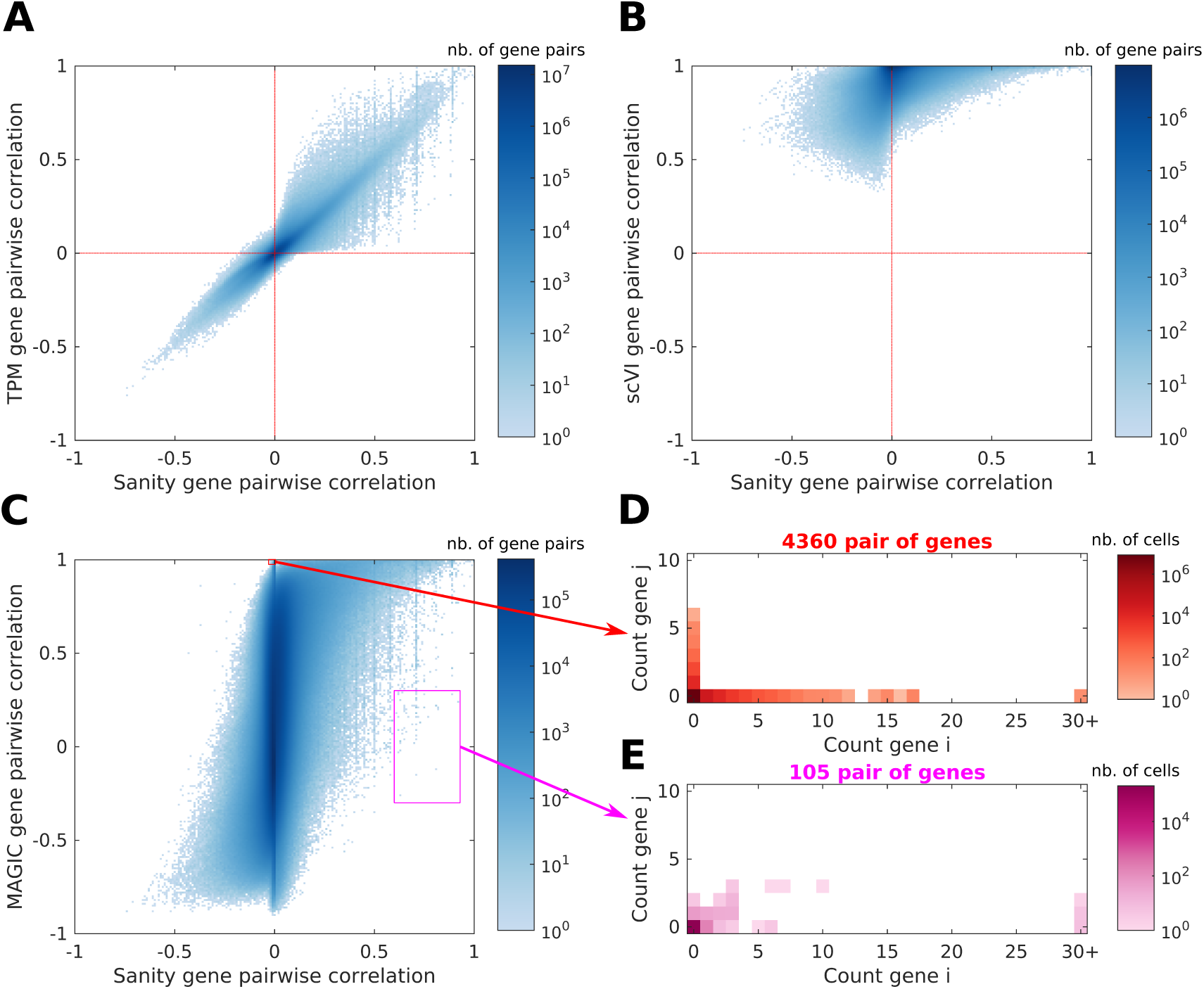
**A**: Density plot of Pearson correlations for all pairs of genes as inferred by *Sanity* (x-axis) against the correlations inferred by *TPM* (y-axis). The color scale shows the density in *log*_10_ number of gene pairs and values log_10_(0) are shown in white. **B**: Density plot as in panel A, but now comparing the correlations inferred by *Sanity* and *scVI*. **C**: Density plot as in panel A but now comparing the correlations inferred by *Sanity* and *MAGIC*. The red and the magenta rectangles indicate the pairs of genes analyzed in panels D and E. The red rectangle contains all pairs of genes with correlation above 0.975 for *MAGIC* and between −0.03 and 0.005 for *Sanity*. The magenta rectangle contains all pairs of genes with correlation between −0.3 and 0.3 for *MAGIC* and between 0.6 and 0.93 for *Sanity*. **D**: 2-dimensional histogram of counts per cell summed over the 4360 pairs of genes from the red rectangle in panel C. The height of the histogram is shown in log_10_ as a color and values log_10_(0) are shown in white. **E**: 2-dimensional histogram of counts per cell summed over the 105 pairs of genes from the magenta rectangle in panel C.

These observations are confirmed by the overall distributions of pairwise correlations that each of the methods predicts on each of the datasets (Suppl. Fig. S4). The results are highly consistent across datasets and show three main behaviors. First, Sanity, TPM, RawCounts, and scImpute have distributions of pairwise correlations that are highly peaked around zero, i.e. these methods predict that most pairs are not co-expressed. Second, instead of a peak at zero, DCA and MAGIC have almost uniform distributions of pairwise correlations. SAVER’s distribution of pairwise correlations is somewhat in between these two behaviors, i.e. a very broad distribution with a moderate peak at zero. Third, scVI’s pairwise correlations are highly peaked near almost perfect correlation of *r* ∼ 1. Notably, even on the simulated dataset that contains no expression correlations at all, MAGIC and DCA also show broad distributions of pairwise correlations, and scVI again predicts almost all pairs to be perfectly co-expressed. This further supports that the correlations that these methods predict are artefactual.

### Sanity outperforms other methods on clustering cells into subtypes

One of the main applications of scRNA-seq is to identify (novel) cell types and this is generally done by clustering single cell gene expression patterns using a measure of pairwise distances between cells. Since the pairwise distances between cells will depend on the normalization method, we expect different methods to differ in their ability to recover subpopulations of cells. For six of our test datasets, the corresponding study explicitly reported an annotation of cell types present in the dataset, that was typically obtained using a combination of automated clustering, analysis of marker gene expression, and hand curation. To test the performance of the different normalization methods on cell type identification we investigated to what extent the reported cell type annotations could be recovered by application of simple clustering algorithms to the normalized gene expression data.

Taking the Zeisel dataset as an example, we first obtained a simplified visual indication of the clustering structure implied by the different methods by applying the popular t-SNE algorithm [26] (with the default 50 principal components and a perplexity equal to the average number of cells per annotated cluster) to the normalized expression values of each method, and colored the cells according to the annotation of [48] (Fig. 5A). Although it is well-known that it is difficult to interpret these visualizations beyond the fact that neighboring cells in the visualization are typically also neighboring in the full gene expression space, the visualization does suggest that there is considerable disparity across the normalization methods. For example, it appears that TPM, DCA, Sanity, and SAVER separate the cell types more reliably than MAGIC, and scVI. Similar qualitative observations can be made on the other datasets (Suppl. Fig. S5 - S9).

**Figure 5:**
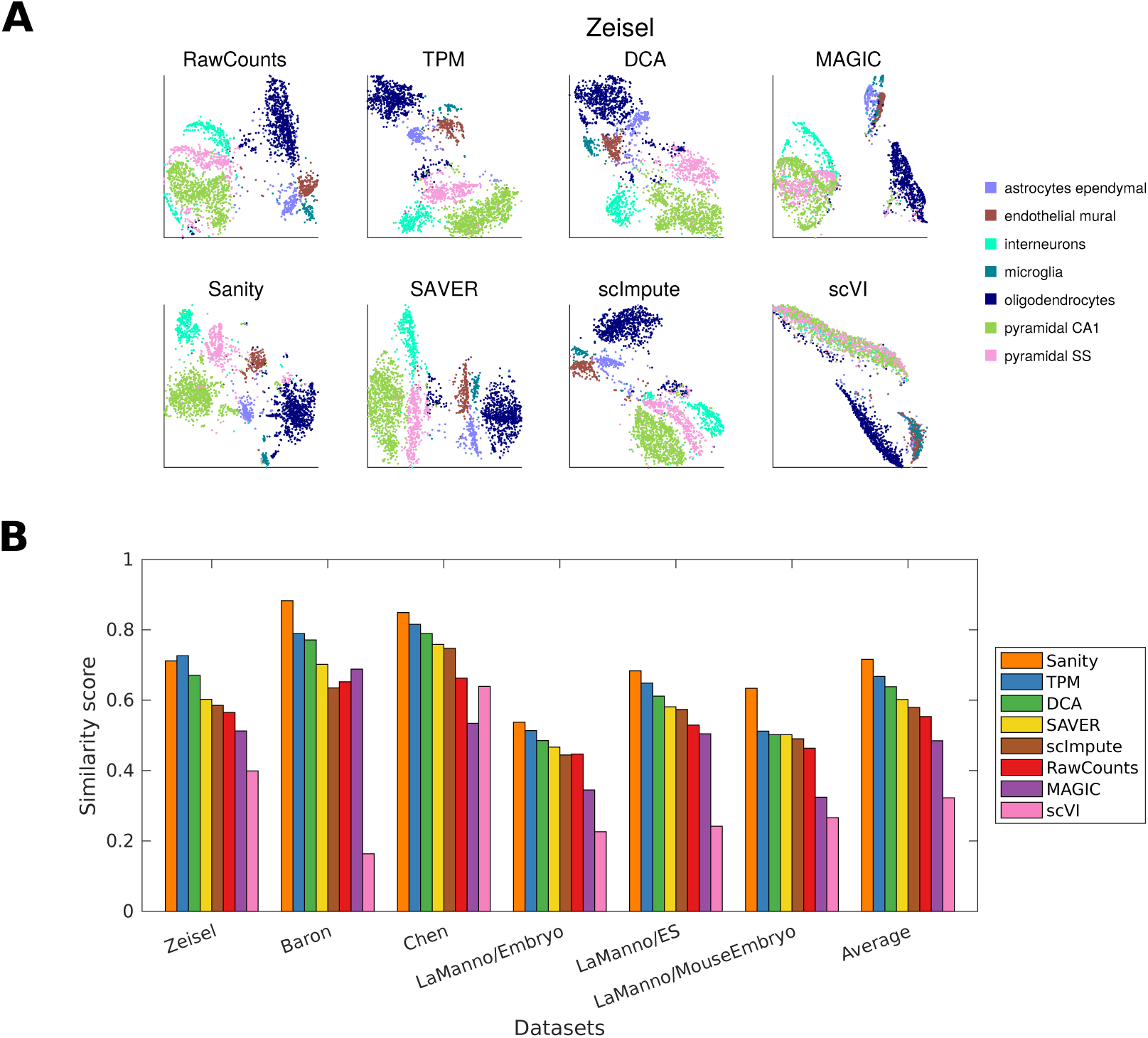
**A**: Each panel shows a t-SNE visualization of the Zeisel dataset using the normalized gene expression values of the method indicated at the top of the panel. Each point represents a cell and is colored according to the cell type annotation given in [48]. **B**: Similarity between the annotated clusters and the clusters inferred by applying hierarchical Ward clustering on the normalized expression values of the different methods. Normalized mutual information, which ranges from 0 (no similarity) to 1 (perfect match) was used as a similarity measure. Each group of bars shows the results for a particular dataset as indicated below it, and the bars are colored according to the normalization method, as indicated in the legend. The last group of bars shows the average similarity per method across all datasets. For ease of viewing, the methods are sorted from left to right according to their average similarity.

To quantify the performance of the different normalization methods we applied, for each dataset and each normalization method, simple hierarchical clustering using Ward’s method [52]. That is, starting with each cell as its own cluster, at each step two clusters are fused so as to minimize the sum of the variances across all clusters. This is iterated until the number of clusters equalled the number of annotated cell types. We then calculated, for each method and dataset, the similarity between the annotated clusters and the inferred clusters using the normalized mutual information as a similarity measure (see Supplementary Methods). As shown in Fig. 5B, Sanity outperforms all other methods on all datasets except the Zeisel dataset, where the TPM method obtains slightly higher similarity with the annotated clusters. The TPM normalization generally has second best performance, followed by DCA and then SAVER. We also note that MAGIC and scVI generally perform least well. Very similar results are observed when using a different similarity metric such as the rand index (Supplementary Methods) or a different clustering algorithm such as k-means (Suppl. Fig. S10).

Thus, although the performance differences are not large, Sanity’s normalized expression estimates generally outperform the other methods in identifying subtypes of cells.

### Sanity outperforms other methods on identification of differentially expressed genes

As a final example of downstream analysis we consider the ability of the normalized expression values to identify genes that are upregulated in particular subtypes of cells. That is, we aim to identify genes whose average expression in a given subtype of cells is significanlty higher than its average in all other cells. A simple and standard statistic for comparing the averages of populations is the *t*-statistic and we used this to identify upregulated genes for each cell type in a given dataset. In particular, for each gene *g* and each cell type *k* annotated in a given dataset, we calculated a *t*-statistic

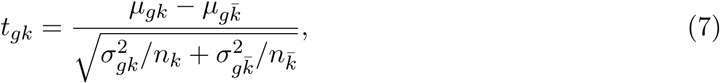

where *μ*_*gk*_ is the average of the normalized expression values of gene *g* in cells of type *k*, 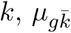 is the average in all other cells, 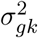 and 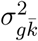 the corresponding variances in normalized expression levels, and *n*_*k*_ and 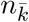 the number of cells in type *k* and the number of all other cells. The *t*-statistic *t*_*gk*_ quantifies that statistical evidence that gene *g*’s average expression in cell type *k* is higher than in the other cells. To predict a set of upregulated genes, one would then pick a cut-off in *t*-statistic corresponding to a particular rate of false discovery (FDR), e.g. a 5% FDR. By applying this procedure to the normalized expression values of each method we derived, for each method, a set of upregulated genes for each cell type *k* of a given dataset of interest.

To test the performance of these predicted sets of upregulated genes we compared these lists with similar lists of predicted upregulated genes from the original publications. For 3 of our test datasets, i.e. the Zeisel and two LaManno datasets, the authors published, for each identified cell type, a list of genes that had higher average expression in the cell type compared to the other types of cells [48, 51]. These lists were obtained using a fairly complex regression procedure and it is of course debatable whether these published lists can be treated as a gold standard. However, since they were obtained using a method that is very different from our simple *t*-statistic, we reasoned that the match to these reference lists can still be used to assess the relative performance of the different normalization methods.

For each normalization method we calculated a precision-recall curve by producing one sorted list of the *t*-statistics *t*_*gc*_ for all genes in all subtypes and, as a function of a cut-off on *t*, compared the predicted set of significantly upregulated genes, with the reference lists published in the original study. Figure 6 shows the precision-recall curves obtained for each of the methods on each of the 3 datasets for which reference lists were available. The colored dots indicate the sensitivity and positive predictive values (PPV) that are obtained for each method when using a *t*-statistic cut-off corresponding to a 5% FDR. We see that, for each dataset, Sanity achieves the highest accuracy of predictions, i.e. a higher PPV at a given sensitivity than all other methods. The simple TPM method achieves the next best performance, MAGIC and scVI perform hardly better than random, and all other methods are somewhere in between. Note that, at a 5% FDR, the more complex DCA, MAGIC, and scVI methods all predict very large numbers of upregulated genes which leads to low PPVs. In summary, these results suggest that Sanity’s normalized expression levels also achieve highest accuracy for downstream identification of differentially regulated genes.

**Figure 6:**
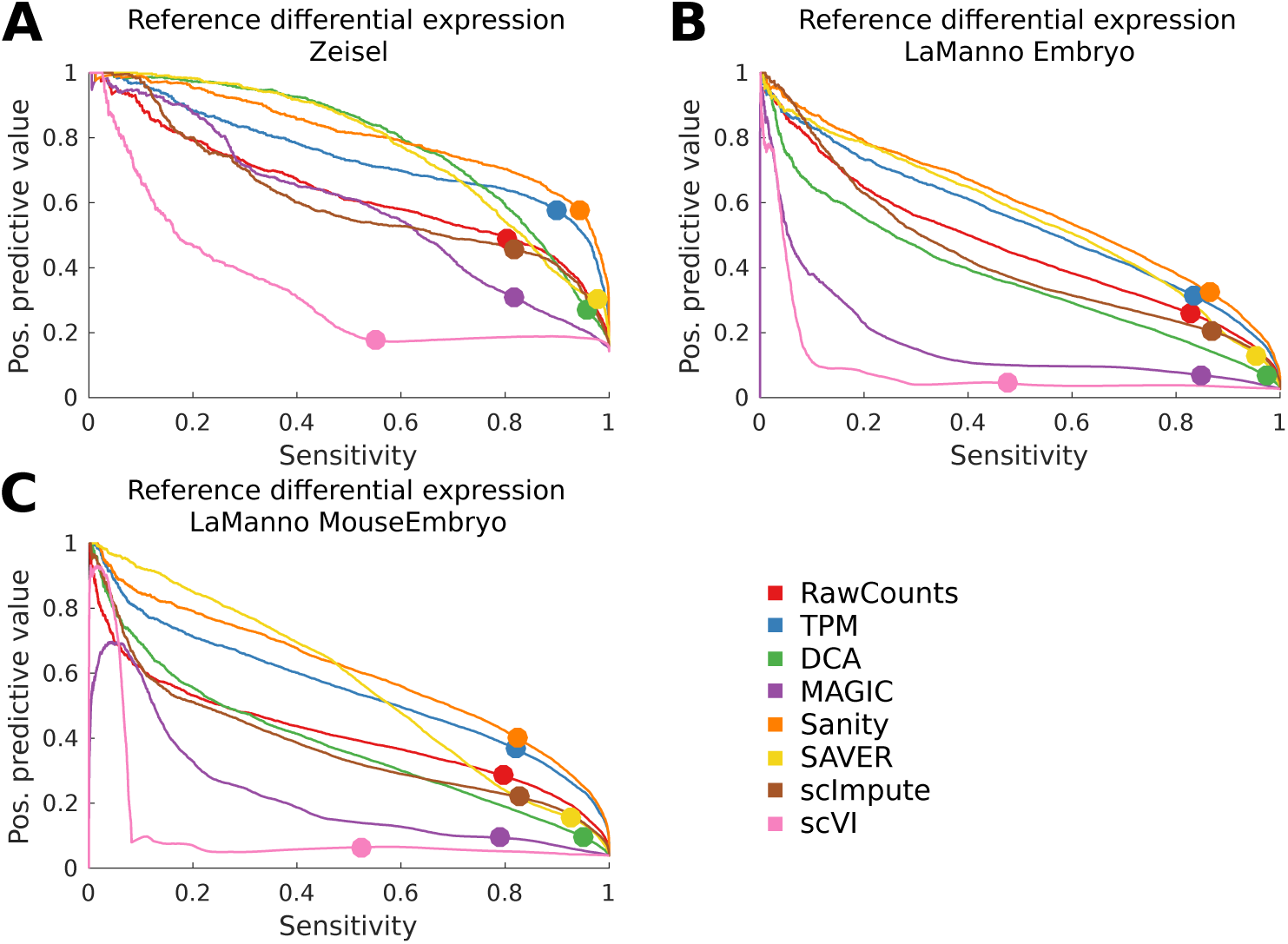
Precision recall curves showing the positive predictive value, i.e. the fraction of predicted upregulated genes that correspond to upregulated genes in the corresponding reference list, as a function of sensitivity, i.e. fraction of all genes in the reference lists that were predicted, as obtained using the *t*-statistics for each of the normalization methods (colors) for the Zeisel (panel **A**) and two LaManno datasets (panels **B** and **C**). The dots show the values that are obtained when using a cut-off on the *t*-statistic corresponding to a false discovery rate of 5%.

### The Sanity software

Sanity was implemented in C and is freely available for download at https://github.com/jmbreda/Sanity. The raw UMI count tables for each of the scRNA-seq datasets, as well as all the normalized expression values as inferred by each of the methods are available from this website as well.

## Discussion

Recent technologies for quantitatively measuring the epigenetic states of single cells are promising to revolutionize our understanding of the mechanisms by which cell fate and identity are regulated in animals, and there has been a surge in the use of these methods. However, even for the most popular scRNA-seq method, there is so far little agreement within the community as to how such single-cell expression data should be processed and normalized. In particular, it has so far been challenging to define a normalization procedure that, on the one hand, deals with the specific artefacts and noise introduced by the scRNA-seq measurement process while, on the other hand, providing quantification of the expression states of cells that have direct biological interpretation. In particular, only when the normalized expression methods provide quantities with a concrete biological interpretation will it be possible to integrate results of scRNA-seq experiments from different protocols employed by different labs with expression data from other experimental techniques. So far, such normalization methods have been lacking.

Here we developed a Bayesian normalization procedure that achieves these objectives, and is derived from first principles using only two basic assumptions. First, we characterize a cell’s gene expression state by the vector of log transcription quotients (LTQs) across genes, i.e. the logarithms of the expected fractions of the transcript pool for each gene. Second, estimating these LTQs within a Bayesian setting requires choosing a prior distribution and we chose to characterize the distribution of LTQs of each gene by just its mean and variance across cells. Given only these two assumptions the entire procedure follows from first principles, deterministically, and without any tunable parameters. Given a table of UMI counts for each gene (or transcript) across cells, our Sanity method returns estimates of LTQs and their error bars across all genes and cells. Importantly, these estimates correct both for the Poisson noise that is inherent in the process of transcription, as well as the sampling noise associated with the scRNA-seq measurements, so that the variance in normalized expression levels across cells reflects changes in rates of transcription and mRNA decay rather than biological or technical sampling noise.

Although our normalization method makes only a minimal number of assumptions, one may ask how arbitrary these assumptions are. If one accepts that biological and technical sampling noise do not reflect changes in gene expression state, that expression changes should be measured in terms of fold-changes rather than absolute changes, and that an overall ‘cell size’ rescaling of the expression levels of all genes by the same amount does not reflect a change in expression state, then LTQs are the most general representation of a cell’s expression state. Similarly, our prior distribution over LTQs for a gene is the least assuming, i.e. maximum entropy, distribution that is consistent with only a given mean and variance. This prior thus also aims to minimize the strength of the assumptions that the method makes. In this sense, we think that our method makes the most conservative assumptions that are consistent with current knowledge. To improve on these assumptions we would have to supply specific biological information to determine more informative priors on the gene expression states that cells can take on.

Our comparison of Sanity with other normalization methods showed that the Sanity’s normalized expression values outperform other methods on basic downstream processing tasks such as clustering cells into subtypes and identifying differentially expressed genes. More importantly, however, we showed that all other methods produce a representation of the data that is severely distorted in one or more respects. Of all alternative methods that we evaluated, the simple TPM method produces the most reasonable representation of the data and it also performs second best on the downstream processing tasks. The main problem with the TPM method is that the variations in normalized expression levels are dominated by Poisson fluctuations. This not only causes there to be a complete lack of correlation between true biological variability of genes and the variability of the normalized expression values, it also causes low expressed genes to be predicted to be most variable, whereas in reality low expressed have least evidence of true variation in gene expression. The simple RawCounts method, and also the scImpute method that produces results highly similar to those of the RawCounts method, both suffer from this same problem, and additionally have the problem of not correcting for variation in total UMI count across cells.

More striking, however, are the severe problems with the normalized expression values produced by the more sophisticated SAVER, DCA, MAGIC, and scVI algorithms. In particular, these methods produce not only strong artefactual correlations of the normalized expression values with the total UMI count in each cell, they also predict very large numbers of co-expressed genes when there is no evidence for co-expression at all in the raw data. The fact that this even occurs on synthetic data where there are no co-expressed genes at all confirms that such spurious correlations are inherently introduced by these normalization procedures.

We believe that these spurious correlations are introduced because all these methods confound noise removal with fitting the data to a lower-dimensional representation. Although it reasonable to assume that the possible states that cells can take on is much lower-dimensional than the full dimensionality of the transcriptome data, the problem of finding such lower-dimensional representations should be clearly distinguished from the problem of correcting for the biological and technical noise. Not only does this noise affect all genes almost independently, but because Poisson sampling noise scales with absolute expression level, different genes are affected by such noise to different extends and this may be erroneously mistaken for ‘structure’ in the data. Indeed, even though methods such as SAVER, DCA, MAGIC, and scVI specifically normalize for the total UMI count per cell, their normalized expression levels show strong correlations (and anti-correlations) with total UMI count. Thus, unless the process of noise removal and normalization is carefully separated from fitting of the data to lower-dimensional representations, artefactual correlations are likely to be introduced.

This is not to say that searching for lower-dimensional representations of the transcriptome data is not an important problem. Indeed, finding biologically meaningful lower-dimensional representations of genome-wide gene expression states is a key challenge in this field. However, we believe that this is a very hard problem in general, and it is currently not clear whether this problem is even solvable in principle, i.e. we are not aware of mathematical results that show under what conditions a lower-dimensional manifold embedded in a very high dimensional space can be reliably reconstructed from a limited number of noisy measurements. Our belief is that, rather than black box procedures for dimensionality reduction, progresss in understanding the genome-wide structure of expression data will crucially depend on connecting transcriptome data to the underlying molecular mechanisms, e.g. the folding of the chromosome, chromatin accessibility at enhancers and promoters, and the binding and unbinding of transcription factors.

However, whatever approach is taken to finding lower-dimensional representations of gene expression states, a prerequisite is that the raw data are first carefully normalized and corrected for both biological and technical sampling noise. The Sanity method that we presented here aims to provide such normalization methodology.

## Supplementary Methods

### Sanity

We denote, for each cell *c* and each gene *g*, the transcription rate a time *t* in the past as *λ*_*gc*_(*t*) and the decay rate of its mRNAs a time *t* in the past as *μ*_*gc*_(*t*). Given these time-dependent transcription and decay rates, we define the *transcription activity a*_*gc*_ of gene *g* in cell *c* as the expected number of mRNAs ⟨*m*_*gc*_⟩ which can be written as the following integral

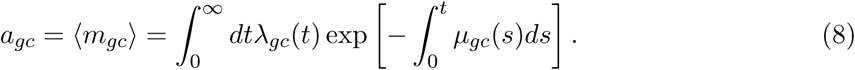

That is, the transcription activity *a*_*gc*_ is a weighted time average of the recent transcription rates a time *t* in the past, with the weight equal to the expected fraction of surviving mRNAs produced at time *t*.

Conditioned on the transcription activity *a*_*gc*_, the distribution of the actual number of mR-NAs *m*_*gc*_ for gene *g* in cell *c* is given by a simple Poisson distribution

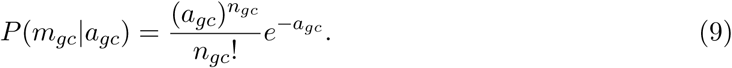

We now assume that, in the scRNA-seq measurement, each mRNA existing in cell *c* has a probability *p*_*c*_ to be captured and sequenced. Given this, the probability that precisely *n*_*gc*_ unique mRNAs will be sequenced for gene *g* in cell *c* is given by

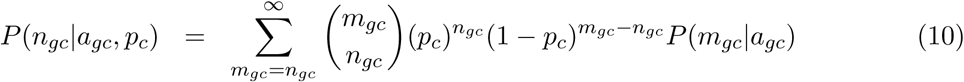

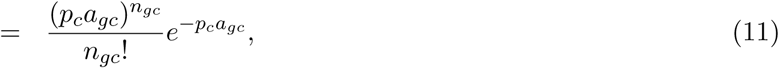

which is still a Poisson distribution.

Next, we define the transcription activity *a*_*gc*_ as a product of the total transcription activity *A*_*c*_ = ∑ _*g*_ *a*_*gc*_ in cell *c* and a *transcription quotient α*_*gc*_:

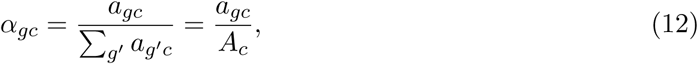

i.e. *α*_*gc*_ is the expected fraction of all mRNAs in cell *c* that are mRNAs for gene *g*. If we also define the cell dependent constant *λ*_*c*_ = *p*_*c*_*A*_*c*_, then we can rewrite this Poisson distribution as

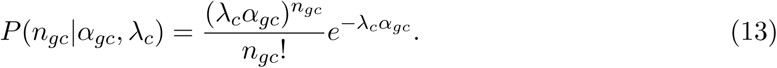

The probability for the entire data-set in cell *c* has the form:

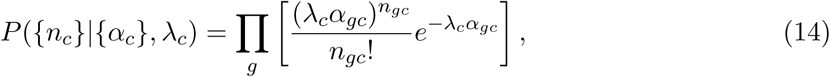

where the notation {*n*_*c*_} refers to all counts *n*_*gc*_ for cell *c*, and {*α*_*c*_} refers to all transcription quotients *α*_*gc*_ for cell *c*. Next, we remove the dependence on the constant *λ*_*c*_ by setting it to its maximum likelihood value. Noting that ∑_*g*_ *α*_*gc*_ = 1 per definition, the dependence of the likelihood *P* ({*n*_*c*_}|{*α*_*c*_}, *λ*_*c*_) on *λ*_*c*_ has the form 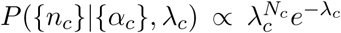, with *N*_*c*_ = ∑ _*g*_ *n*_*gc*_ the total number of sequenced mRNAs in cell *c*. Thus, the value of *λ*_*c*_ that maximizes *P* ({*n*_*c*_}|{*α*_*c*_}, *λ*_*c*_) is simply *λ*_*c*_ = *N*_*c*_. Substituting this we obtain

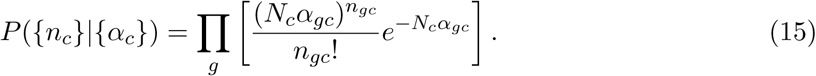

That is, the number of sequenced mRNAs *n*_*gc*_ for each each gene *g* in cell *c* is still a Poisson distribution with expectation value *N*_*c*_*α*_*gc*_. The probability of the entire dataset of counts {*n*} given all transcription quotients {*α*} is given by simply taking the product of this expression over all cells, i.e

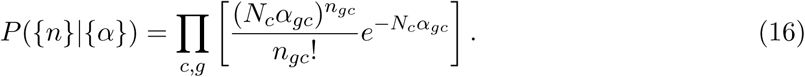

Instead of trying to estimate the *α*_*gc*_ for all genes at once, we will focus on one specific gene *g* at a time, and infer how *α*_*gc*_ varies across the cells *c*. Note that if we collect all the terms that depend on the *α*_*gc*_ of single gene *g* we obtain

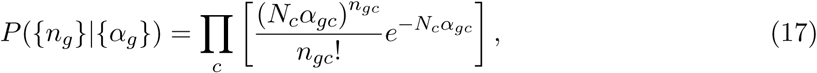

where {*n*_*g*_} is the set of counts for gene *g* and {*α*_*g*_} is the set of transcription quotients for gene *g*.

Finally, without loss of generality, we will write the transcription quotients *α*_*gc*_ in terms of the average quotient of the gene *α*_*g*_ and a log-fold change *δ*_*gc*_ in a given cell *c*, i.e.

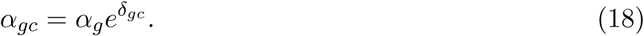

In terms of these parameters we have

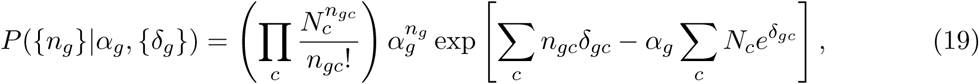

where *n*_*g*_ is the total number of sequenced mRNAs for gene *g*.

### Marginalizing over the average transcription quotient *α*_*g*_

We now first focus on estimating the log fold-changes *δ*_*gc*_. We return to estimating the overall average transcription quotient *α*_*g*_ once we have determined these. To marginalize expression (19) over *α*_*g*_ we use a simple uniform prior *P* (*α*_*g*_)*dα*_*g*_ ∝ *dα*_*g*_. Integrating with this uniform prior from 0 to ∞ we obtain

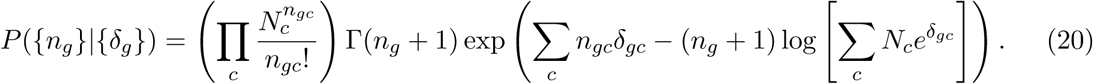

Note that, because *α*_*g*_ is a fraction, we should have really only integrated from 0 to 1, but as long as each gene is only responsible for a small fraction of all UMIs in the cell, the only contribution to the integral comes from values of *α*_*g*_ much smaller than 1, and we can extend the range of the integral to infinity without loss of accuracy. Note also that the factor 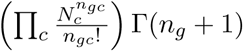 is determined entirely by the counts and does not depend on the *δ*_*gc*_, and we will neglect this prefactor from now on.

### Including prior probabilities for the *δ*_*gc*_

We next introduce prior probabilities over the log fold-changes *δ*_*gc*_. Assuming only that the *δ*_*gc*_ for gene *g* have a variance *v*_*g*_ and mean zero, we use the maximum entropy distribution consistent with these constraints, which is a Gaussian

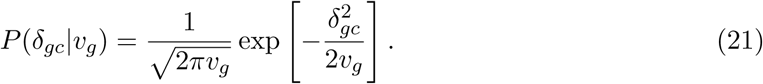

Thus, the prior over the full set {*δ*_*g*_} of log fold-changes for the *C* cells is given by

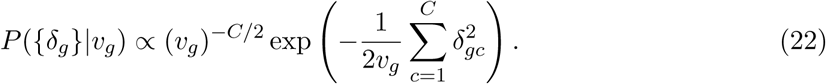

Combining the prior with the likelihood we obtain

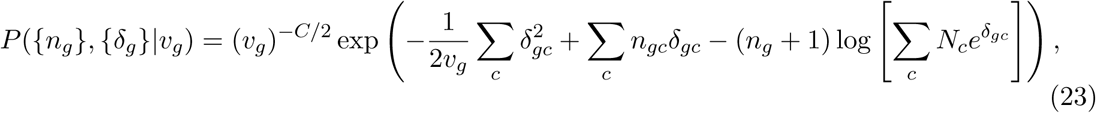

up to a prefactor that does not depend on the parameters *δ*_*gc*_ and *v*_*g*_.

### Calculating *P* ({*n*}|*v*_*g*_) using the Laplace approximation

We next focus on calculating the probability *P* ({*n*_*g*_}|*v*_*g*_) of the data given only the variance *v*_*g*_. To obtain the probability *P* ({*n*_*g*_}|*v*_*g*_), we need to integrate over all possible *δ*_*gc*_. As the integral is close to Gaussian in form, we will assume we can approximate the integral by the Laplace approximation, i.e. by approximating the log-likelihood *L*({*δ*_*g*_}, *v*_*g*_) = log [*P* ({*n*_*g*_}, {*δ*_*g*_}|*v*_*g*_)] by expanding it to second order around its maximum. The log-likelihood has the form

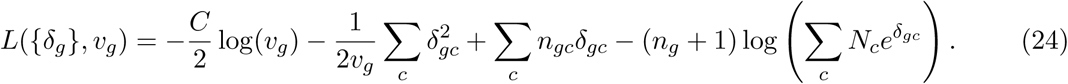

Taking derivatives with respect to the *δ*_*gc*_, the equations for the optimum become

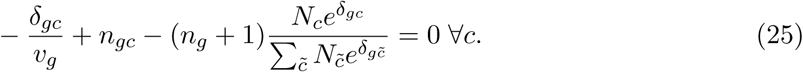

To solve this equation we are going to multiply the equation by *v*_*g*_ and then define the *c*-independent quantity

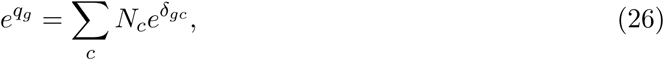

the normalized quantities

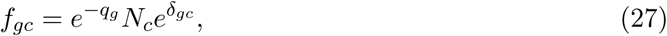

which sum to 1, i.e. ∑ _*c*_ *f*_*gc*_ = 1, and the *c*-dependent quantities

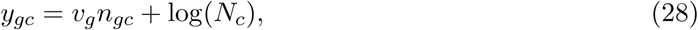

which are directly determined by *v*_*g*_ and the data.

In terms of these quantities the equations for the optimum become

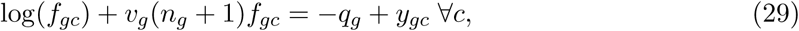

whose solution is

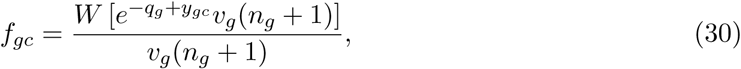

with *W* (*x*) the Lambert W-function (also called productlog). Note, however, that the solution depends on *q*_*g*_, which itself depends on the *f*_*gc*_. However, since ∑ _*c*_ *f*_*gc*_ = 1 per definition, we can sum equation (30) over *c* to obtain the following consistency equation for *q*_*g*_

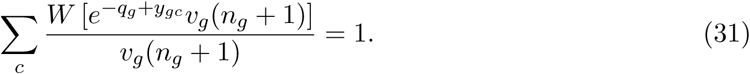

In the above equation, everything is determined either by the data (*n*_*gc*_, *n*_*g*_, and *N*_*c*_) or the variance *v*_*g*_, except for the unknown constant *q*_*g*_, which needs to be solved for numerically. We can perform a binary search to find the value of *q*_*g*_ for which equation (31) is satisfied. Note also that the expression on the left hand side of equation (31) is a monotonically decreasing function of *q*_*g*_, guaranteeing that there is only a single solution for *q*_*g*_.

Once *q*_*g*_ has been determined, we obtain the *f*_*gc*_ from equation (30) and we obtain the optimal 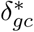 as

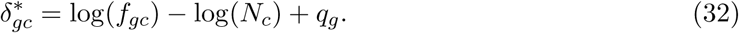

Note that these optimal 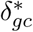 are functions of the variance *v*_*g*_, which we from now on will express explicitly in our notation.

Substituting the optimal 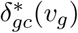 into equation (24) we obtain the optimal log-likelihood *L*_*_(*v*_*g*_). By expanding the log-likelihood to second order around its maximum, the probability *P* ({*n*_*g*_}, {*δ*_*g*_}|*v*_*g*_) can then be rewritten as

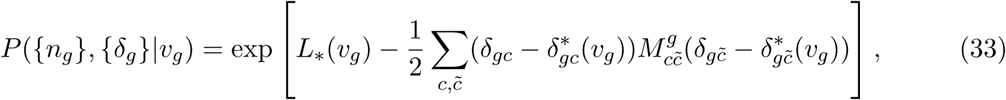

where the matrix *M* ^*g*^ is given by the second derivatives of the log-likelihood around its optimum, i.e.

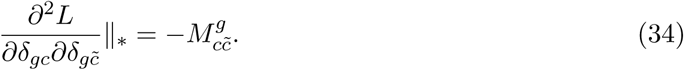

We find

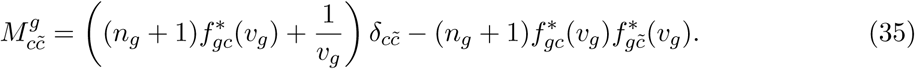

The integral over the likelihood can now be easily written in terms of the determinant of the matrix *M* ^*g*^, given us for the marginal probability of the data as a function of *v*_*g*_:

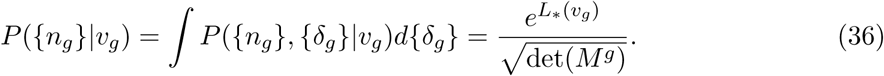

Finally, given the relatively simple structure of the matrix *M* ^*g*^, we use the *matrix determinant lemma* and write the determinant as

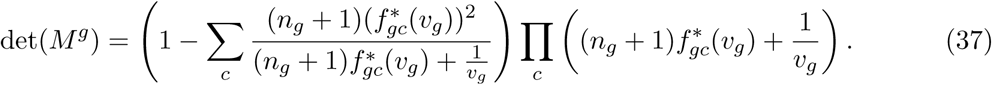

### Posterior *P* (*v*_*g*_|{*n*}) over variance *v*_*g*_

To obtain a posterior over the variance *v*_*g*_ we need a prior over the variance *v*_*g*_, for which we will use a scale prior, i.e. uniform in the logarithm of *v*_*g*_: *P* (*v*_*g*_)*dv*_*g*_ ∝ *d* log(*v*_*g*_). Note, however, that our solution of *P* ({*n*}|*v*_*g*_) involved a numerical determination of *q*_*g*_, so that we do not have an analytical formula for *P* (*v*_*g*_|{*n*}). In order to approximate the full posterior *P* (*v*_*g*_|{*n*}) we pick a range [*v*_min_, *v*_max_] within which we presume all *v*_*g*_ fall, divide this range into *B* bins of equal size in log(*v*_*g*_), and calculate *P* ({*n*}|*v*_*g*_) for each bin *b*. Per default we choose [*v*_min_, *v*_max_] = [0.01, 20] since this covers the range of observed variances in the datasets we considered. Trading off speed versus accuracy we chose *B* = 116 bins by default, so that the variance increase by about 5% from one bin to the next. However, if desired these values can be changed by the user.

Let *v*_*b*_ denote the variance of bin *b* and *L*_*b*_ the log-likelihood log[*P* ({*n*}|*v*_*b*_)]. We then approximate the full posterior *P* (*v*_*b*_|{*n*}) by a distribution over a finite number of points:

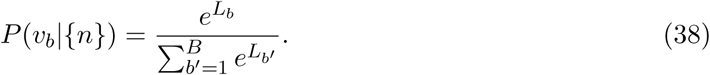

### The posterior *P* ({*δ*_*g*_}|{*n*}, *v*_*g*_) of log-fold changes given a variance *v*_*g*_

For a given value of the variance *v*_*g*_, the posterior distribution over the log fold-changes *δ*_*gc*_ is given by a multi-variate Gaussian with means 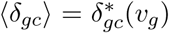 and a covariance matrix *C* given by the inverse of the matrix *M*^g^. In particular, the variances var(*δ*_*gc*_) of the log fold-changes across cells are given by the diagonal elements of the inverse of *M* ^*g*^. Fortunately, given the relatively simple structure of the matrix *M* ^*g*^, we can also obtain analytical expressions for these variances. In particular, the components (*c, c*) of the inverse of *M* ^*g*^ are given by the ratio of the minor [*M* ^*g*^]_(*c,c*)_ (the determinant of matrix *M* ^*g*^ with the *c*th row and column removed) and the determinant of the full matrix. We have

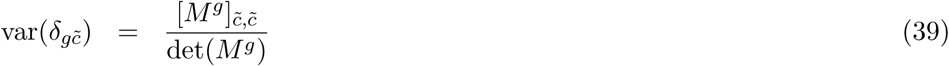

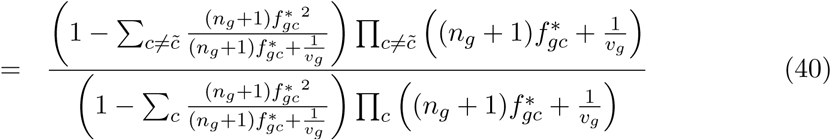

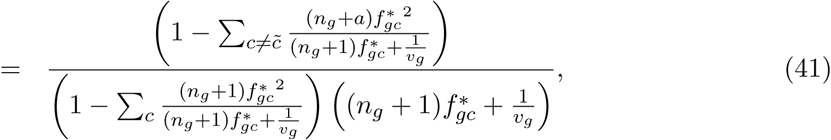

where again it should be noted that the 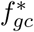 are themselves functions of *v*_*g*_.

A technical complication arises in estimating the variance var(*δ*_*gc*_) when the observed number of UMIs is zero. That is, when *n*_*gc*_ = 0 the log-likelihood *L*({*δ*_*g*_}, *v*_*g*_) can be a highly asymmetric function of *δ*_*gc*_ around its maximum 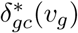. In particular, whereas the fact that no UMIs were observed, i.e. *n*_*gc*_ = 0, ensures that the log-likelihood decreases quickly as *δ*_*gc*_ increases above 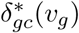, it drops only slowly with decreasing *δ*_*gc*_. That is, when no UMI are observed, we can give a reasonably tight upper bound on *δ*_*gc*_, but *n*_*gc*_ = 0 is consistent with very low *δ*_*gc*_. This asymmetry causes the variance var(*δ*_*gc*_) to significantly overestimate the error-bar in *δ*_*gc*_ toward larger values of *δ*_*gc*_. To fix this problem, we directly set var(*δ*_*gc*_) from its upper bound for cases with *n*_*gc*_ = 0. In particular, note that for a Gaussian distribution with mean *µ* and variance *σ*^2^, the difference between the log-likelihood at the optimum *µ* and at *µ* + *σ* is *L*(*µ*) − *L*(*µ* + *σ*) = (*µ* + *σ* − *µ*)^2^*/*(2*σ*^2^) = 1*/*2. We thus define the 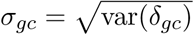 such that the difference between the log-likelihood at 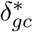 and 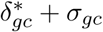 is 1*/*2, i.e. the solution of

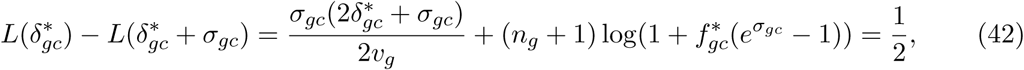

which we determine numerically.

### Final estimates ⟨*δ*_*gc*_⟩ and error-bars *ϵ*_*gc*_

For each value of *v*_*g*_, we have determined the posterior probability *P* (*v*_*g*_|{*n*}) and given a variance *v*_*g*_, we have a Gaussian posterior distribution *P* ({*δ*_*g*_}|{*n*}, *v*_*g*_) over the log fold-changes, with means 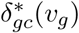 and variances var(*δ*_*gc*_)(*v*_*g*_). Using these, we can now calculate final estimates of the fold changes *δ*_*gc*_. In particular, the expectation value ⟨*δ*_*gc*_⟩ is given by the integral

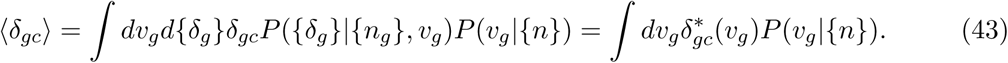

Similarly, we find for the overall error-bar 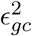

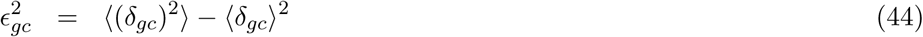

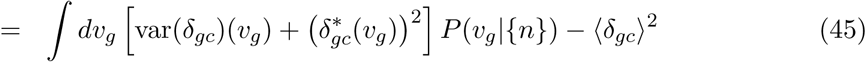

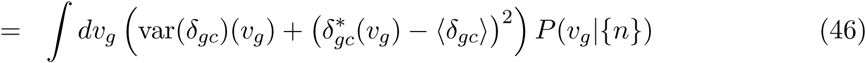

Sanity returns, for each gene *g* in each cell *c*, both the estimated log fold-change ⟨*δ*_*gc*_⟩ and its error-bar *ϵ*_*gc*_.

### Mean expression ⟨log(*α*_*g*_) ⟩

Once we have fitted a set of 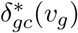 for each *v*_*g*_, and determined the posterior *P* (*v*_*g*_|{*n*_*g*_}) we can now easily estimate the mean log quotient *µ*_*g*_ = log(*α*_*g*_) of each gene. Returning to equation (19), and marginalizing over the *δ*_*gc*_ we find that the posterior over *α*_*g*_ is proportional to the expression (19) in which the *δ*_*gc*_ have been set to 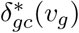:

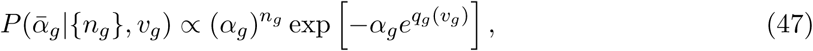

where *n*_*g*_ is the total number of UMIs captured for gene *g*, 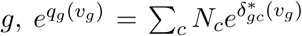 as defined above, and we have explicitly indicated that *q*_*g*_ is a function of the variance *v*_*g*_.

Using (47) the expectation value of log(*α*_*g*_) at a given value of the variance *v*_*g*_ is given by

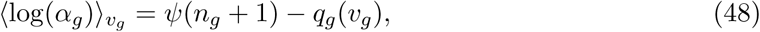

where *ψ*(*x*) is the digamma function, i.e. the derivative of the logarithm of the gamma function. Note also that, since *n*_*g*_ is an integer, we have *ψ*(*n*_*g*_ + 1) is simply related to the Harmonic numbers, i.e. 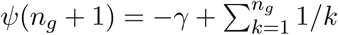, with *γ* ≈ 0.577 the Euler–Mascheroni constant.

To get a final estimate *µ*_*g*_ = (log(*α*_*g*_)) we obtain the weighted average over the variance *v*_*g*_, i.e.

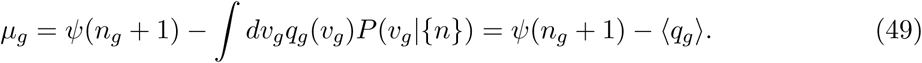

### Error bar on mean expression

Going back to equation (47) we find that the variance in log(*α*_*g*_), at a given value of the variance *v*_*g*_, is given by the derivative of the digamma function:

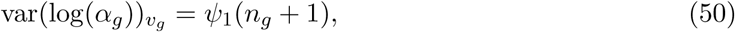

with *ψ*_1_(*x*) the derivative of the digamma function, which is also called the trigamma function. Note that this variance is independent of *v*_*g*_.

The final error-bar *δµ*_*g*_ for log(*α*_*g*_ + 1) is then given

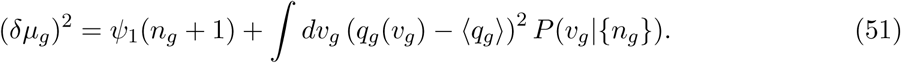

Note that, as for the calculation of the log fold-changes, these integrals over *v*_*g*_ are approximated by sums over the same set of *B* bins.

### Simulated dataset

Defining *N*_*gene*_ the number of genes, *N*_*cell*_ the number of cells, *µ*_*g*_ the mean LTQ of gene *g, v*_*g*_ the variance of the LTQs of gene *g* across cells, *N*_*c*_ the total number of sequenced mRNAs in cell *c, a*_*gc*_ the transcription activity of gene *g* in cell *c*, and *α*_*gc*_ the transcription quotient of gene *g* in cell *c*, we simulated the UMI counts *n*_*gc*_ as follows:

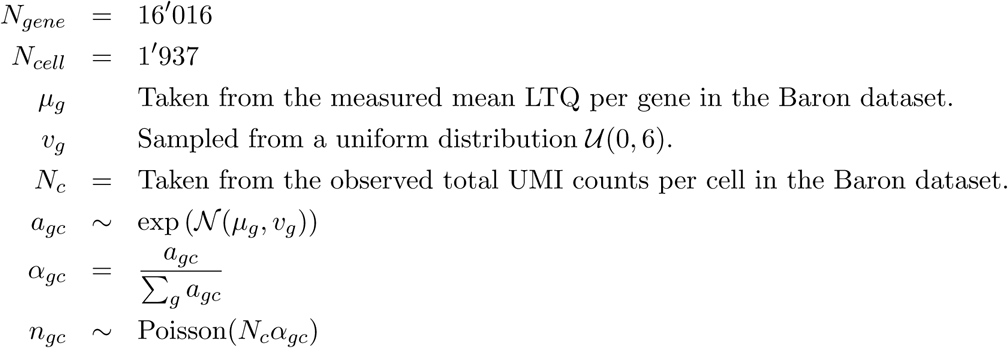

That is, the mean LTQs *µ*_*g*_ were taken from the mean LTQs measured on the Baron dataset. The variances in LTQs were drawn from a Uniform distribution between 0 and 6. The total number of UMIs per cell was chosen identical to the total number of UMIs per cell observed in the Baron dataset. For each gene *g*, the transcription activity *a*_*gc*_ of each cell *c* was then drawn from a log-normal with mean of log(*a*_*gc*_) equal to *µ*_*g*_ and variance *v*_*g*_, and the transcription quotients *α*_*gc*_ were set by normalizing to the total transcription activity of each cell. Finally, the observed UMI counts *n*_*gc*_ were drawn from a Poisson with mean *N*_*c*_*α*_*gc*_ for each gene *g* in each cell *c*.

Figure S11 shows the distributions of the total number of mRNAs per cell, the total number of mRNAs per gene, and the variance in observed mRNA counts for both the Baron dataset and the simulated data. Note that the distributions are highly similarly except for the variances, which are more widely distributed in the simulated data.

### Clustering index

Let the sets {*A*} and {*B*} denote two cell classifications where *A*_*i*_ ∈ ℕ_+_ and *B*_*i*_ ∈ ℕ_+_ denote the class numbers of cell *i* in the two classifications, and *i* = 1, …, *C*, with *C* the number of cells (*i.e.* the number of elements in sets {*A*} and {*B*}).

The distributions and the joint distribution of the two classifications are defined as the frequencies

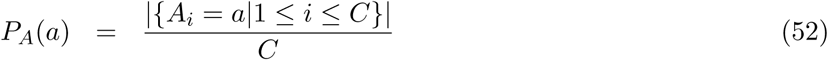

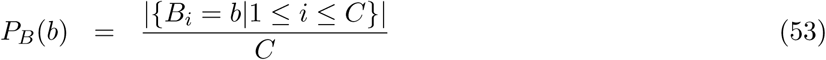

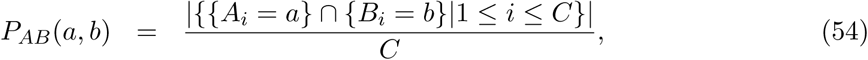

where | · | denotes the cardinality of a set.

The entropy of the distributions and the joint distribution are defined

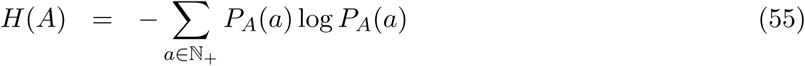

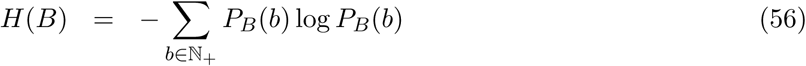

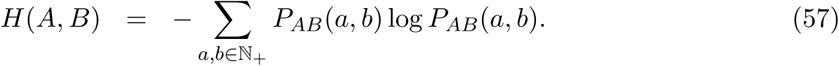

The *mutual information* is defined

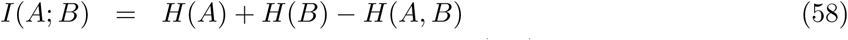

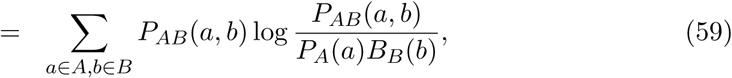

representing the difference between the summed entropy of the 2 distributions and the entropy of the joint distribution.

We compute the *Normalized mutual information* as

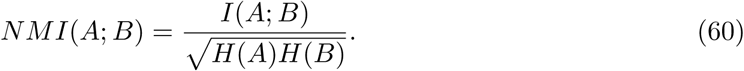

Alternatively, the confusion matrix being defined

**Table S1:** Confusion matrix.

|  | Inferred classes |  |
| --- | --- | --- |
| Reference | $B_i = B_j$ | $B_i \neq B_j$ |
| $A_i = A_j$ | TP | FN |
| $A_i \neq A_j$ | FP | TN |

The *Adjusted rand index* is defined

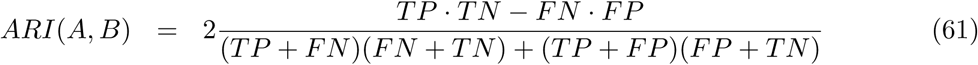

### Differential expression

Let *e*_*gc*_ the log-expression of gene *g* in cell *c, C* an ensemble of cells, and 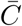 all other cells in the dataset. The *t*-statistic *t*_*gC*_ quantifies the statistical evidence that the average expression of gene *g* in the set *C* differs from the average in all other cells:

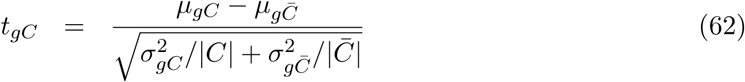

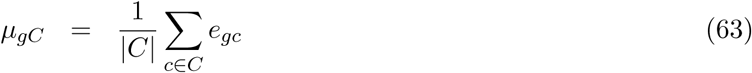

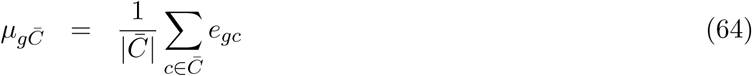

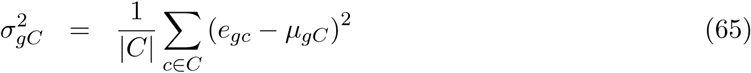

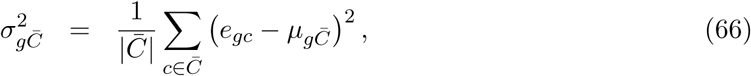

with |*C*| and 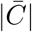 the number of cells in set *C* and its complement, respectively.

Given a *t*-statistic *t*_*gC*_, the *p*-value under a one-side *t*-test that the gene is over-expressed in set *C* is given by

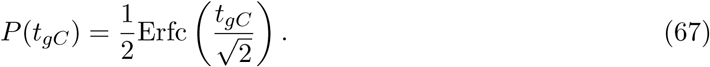

Sorting all genes by the *t*-statistic *t*_*gC*_ the list of over-expressed genes at a false discovery rate of *f* is obtained by picking a cut-off *t*_*c*_ such that average of *P* (*t*_*gC*_) for all genes with *t*_*gC*_ > *t*_*c*_ is *f*.

The reference sets of differentially expressed genes are constructed using a negative binomial generalized linear regression to obtain posterior probability distributions for the class-specific contributions to each gene’s expression (also considering contribution of age and sex and a basal expression per gene) (see [48], Supplementary Materials, Gene expression enrichment analysis).

### Correlation matrix distance

Given 2 correlation matrix *R*_1_ and *R*_2_, the *correlation matrix distance* (CMD) [53] measures a distance between *R*_1_ and *R*_2_, bound between 0 (equal) and 1 (most different) and is defined as

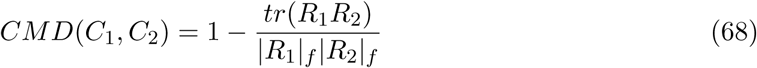

where | · |_*f*_ denotes the frobenius norm, and *tr*(·) the trace.

## Supplementary figures

**Figure S1:**
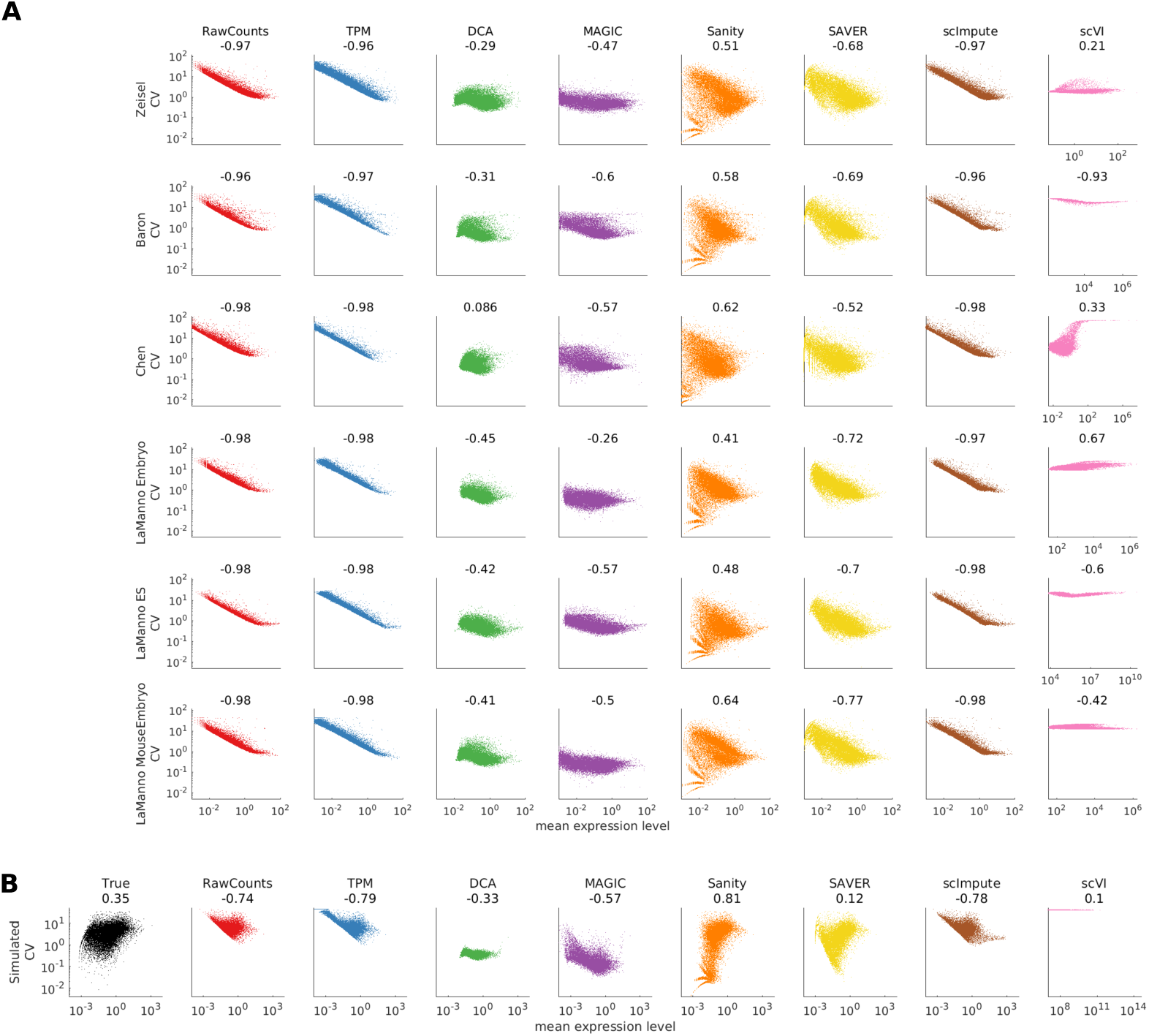
**A** Scatter plots of CV against mean gene expression level. Rows correspond to different scRNA-seq data. Colors and columns correspond to the different methods used to normalize the data.The axis are kept similar across panels, except for *scVI* for which the x-axis is different as the mean expression is on a very different scale compared to the others. **B** Scatter plots of CV against mean gene expression level. The panels and colors correspond to different methods used to normalize the data. The different methods and the correlation between the inferred CV and the true CV is shown on top of each panel.The axis are kept similar across panels, except for *scVI* for which the x-axis is different as the mean expression is on a very different scale compared to the others.

**Figure S2:**
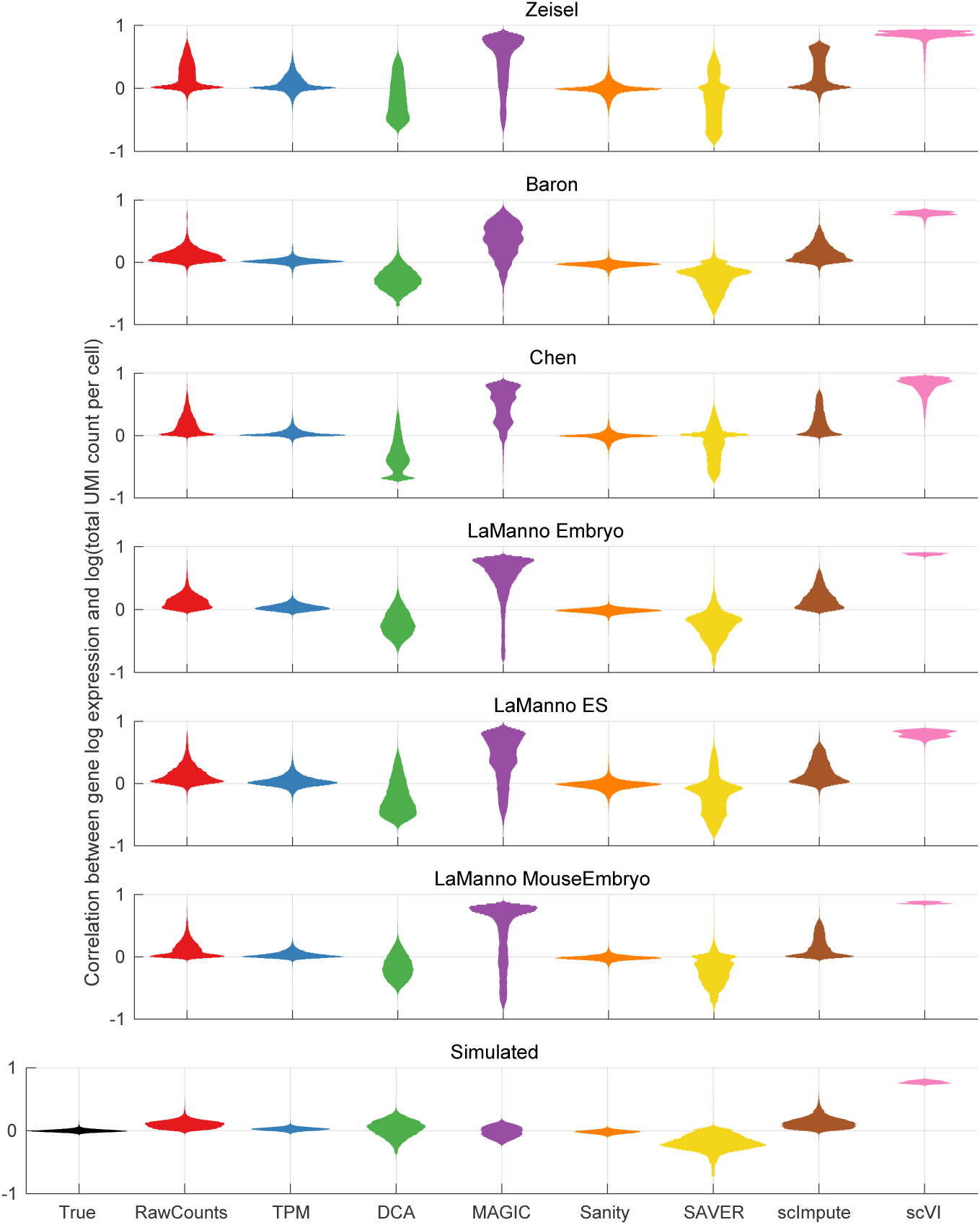
Violin plots of the distribution of correlation coefficients between inferred log expression level of genes and and log of total mRNA molecule count per cell. Rows correspond to different datasets, as indicated on top of each panel. Columns correspond the different methods, as indicated on the x-axis.

**Figure S3:**
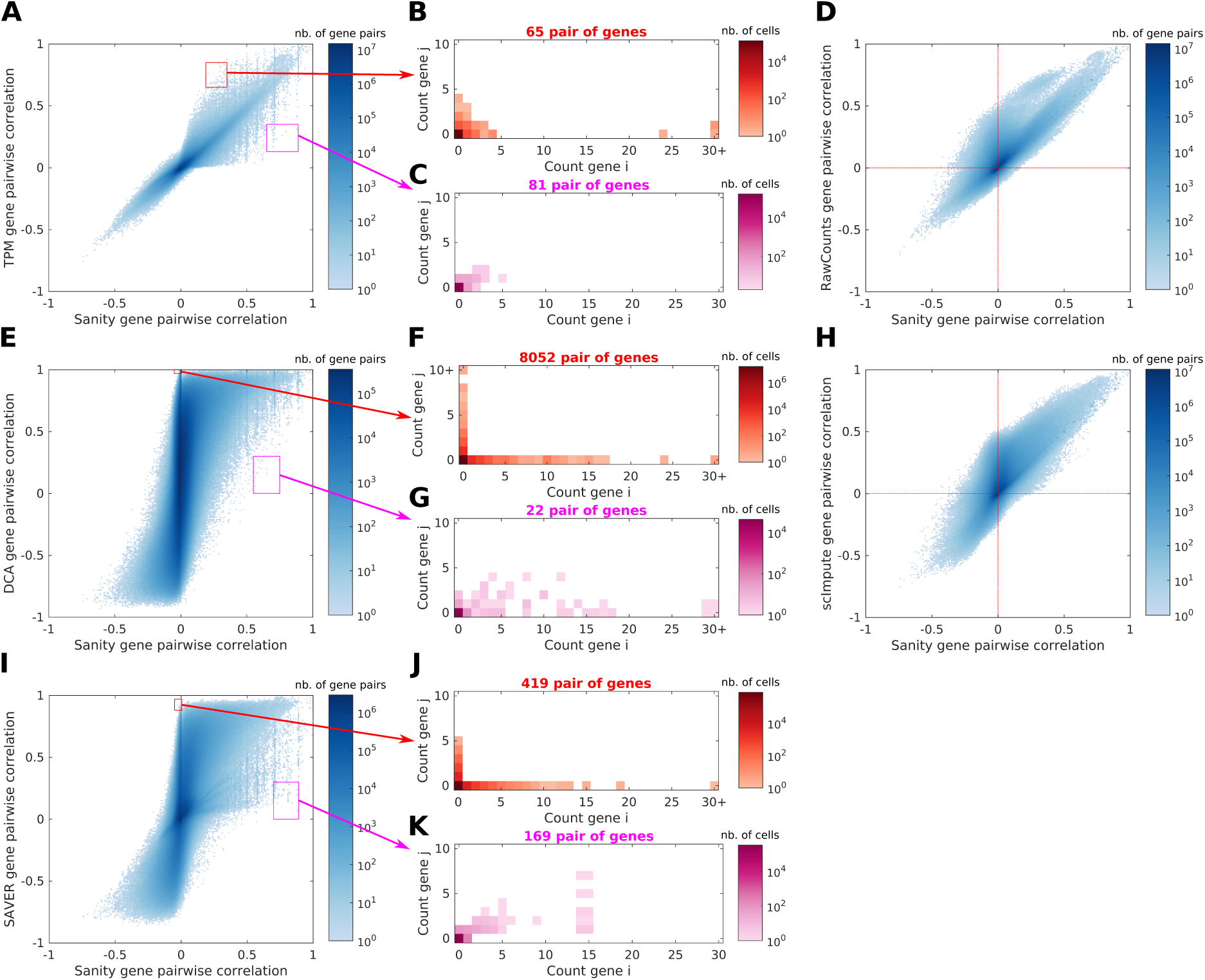
Density plot of the Pearson correlations of normalized expression values of all pairs of genes as inferred by Sanity (x-axis) and the correponding correlation as inferred by TPM (**A**), RawCounts (**D**), DCA (panel **E**), scImpute **H**, and SAVER (**I**)) on the y-axis for the Baron dataset. The color scale shows the density in *log*_10_ of gene pairs and values log_10_(0) are shown in white. For panels **A, E**, and **I**, the red and magenta rectangles show selections of gene pairs for which the two methods disagree most strongly on the correlation. For each such set of pairs, we counted the number of *n*_*i,j*_ across all pairs and all cells for which *i* UMI were observed for the first gene and *j* UMI for the second gene. The panels **B, C, F, G, J**, and **K** show the corresponding 2-dimensional histograms *n*_*i,j*_ for each selected set with the number of pairs indicated above the panel. The height of the histogram is shown in log_10_ as a color and values log_10_(0) are shown in white.

**Figure S4:**
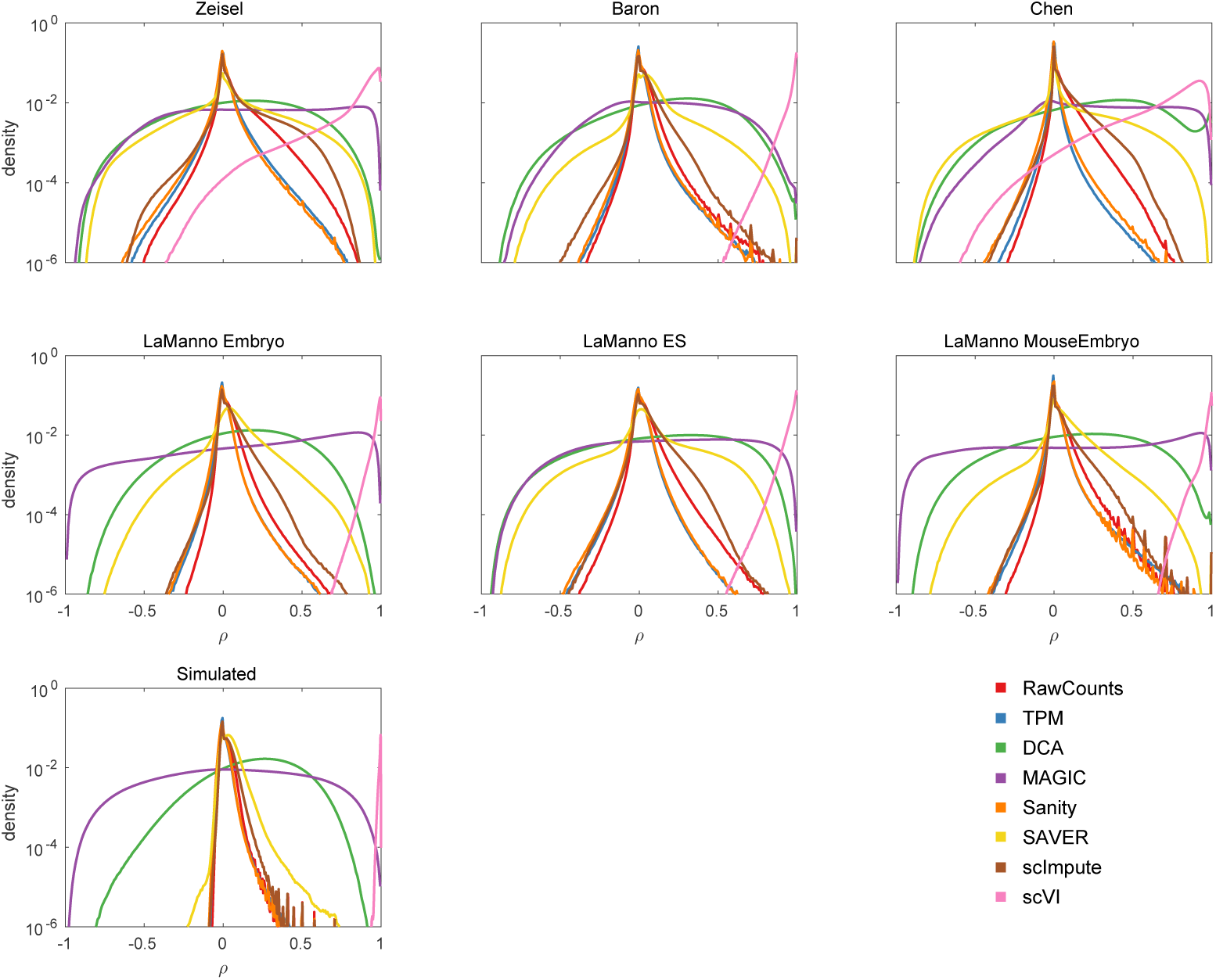
Distributions of the Pearson correlations of all pairs of genes, as inferred by each normalization method. Each panel corresponds to one dataset (indicated at the top of the panel) and each color corresponds to one of the normalization methods, as indicated in the legend. Note that the y-axis is shown on a logarithmic scale.

**Figure S5:**
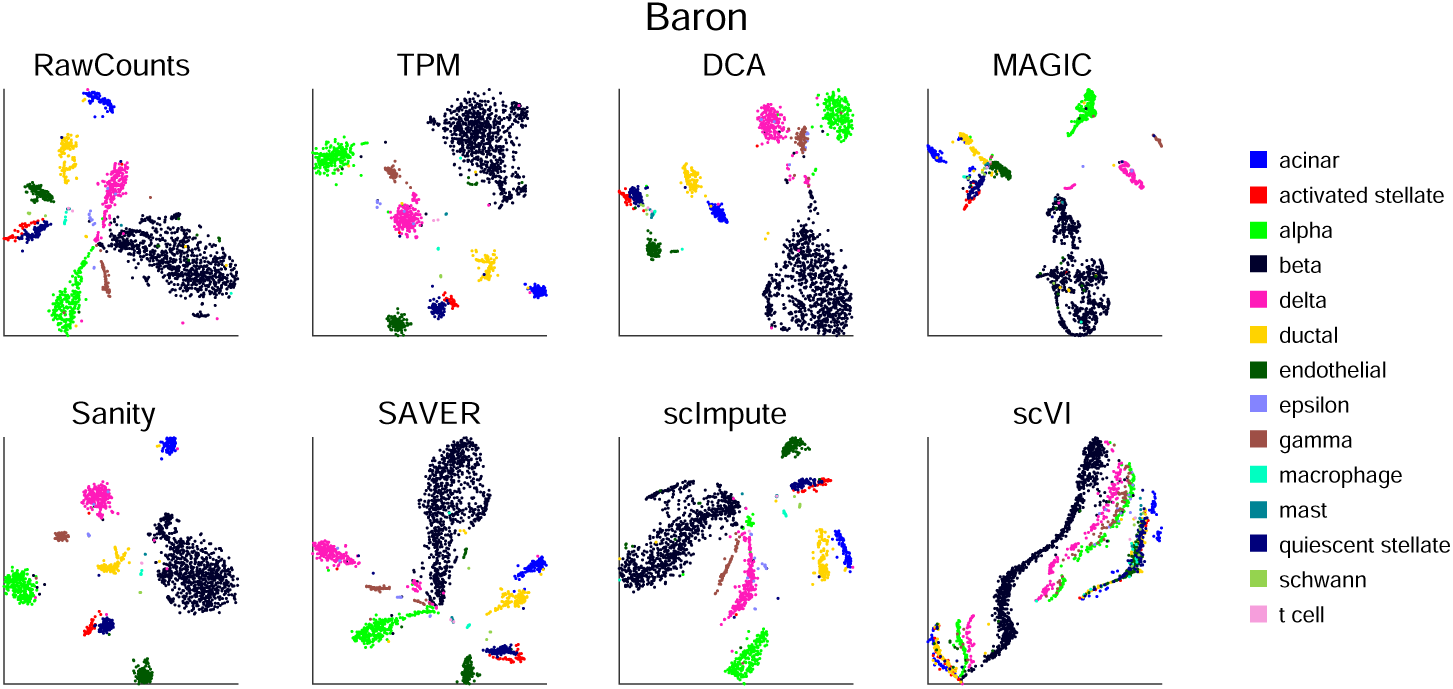
Each panel show a t-SNE visualization of the Baron dataset using the normalized gene expression values of the method indicated at the top. Each point represent a cell and is colored by the cell type annotated in the original publication.

**Figure S6:**
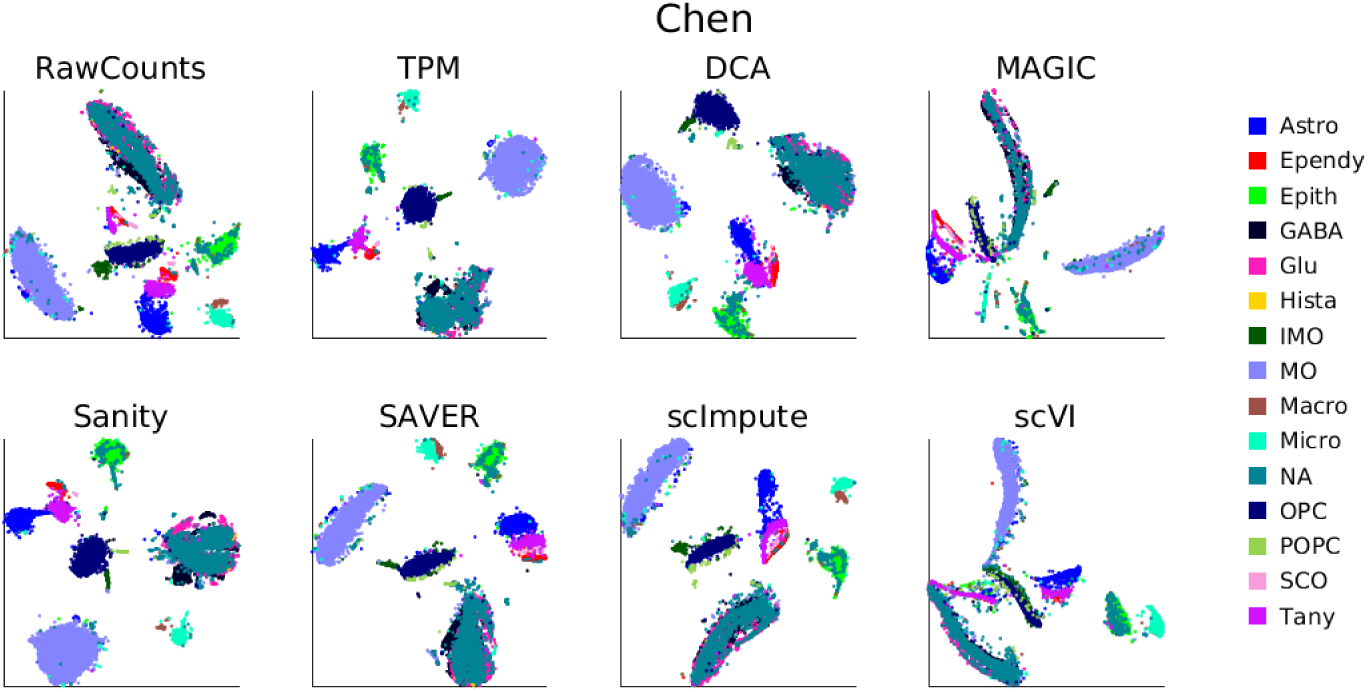
Each panel show a t-SNE visualization of the Chen dataset using the normalized gene expression values of the method indicated at the top. Each point represent a cell and is colored by the cell type annotated in the original publication.

**Figure S7:**
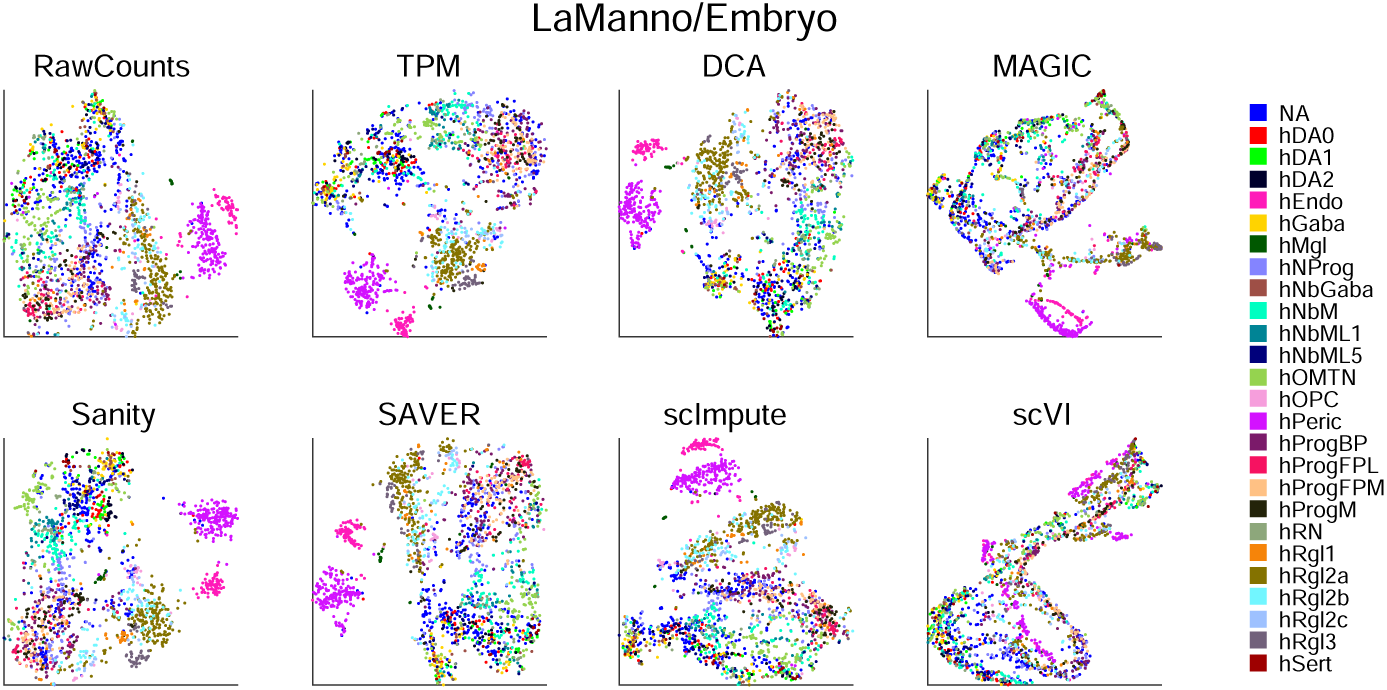
Each panel show a t-SNE visualization of the LaManno/Embryo dataset using the normalized gene expression values of the method indicated at the top. Each point represent a cell and is colored by the cell type annotated in the original publication.

**Figure S8:**
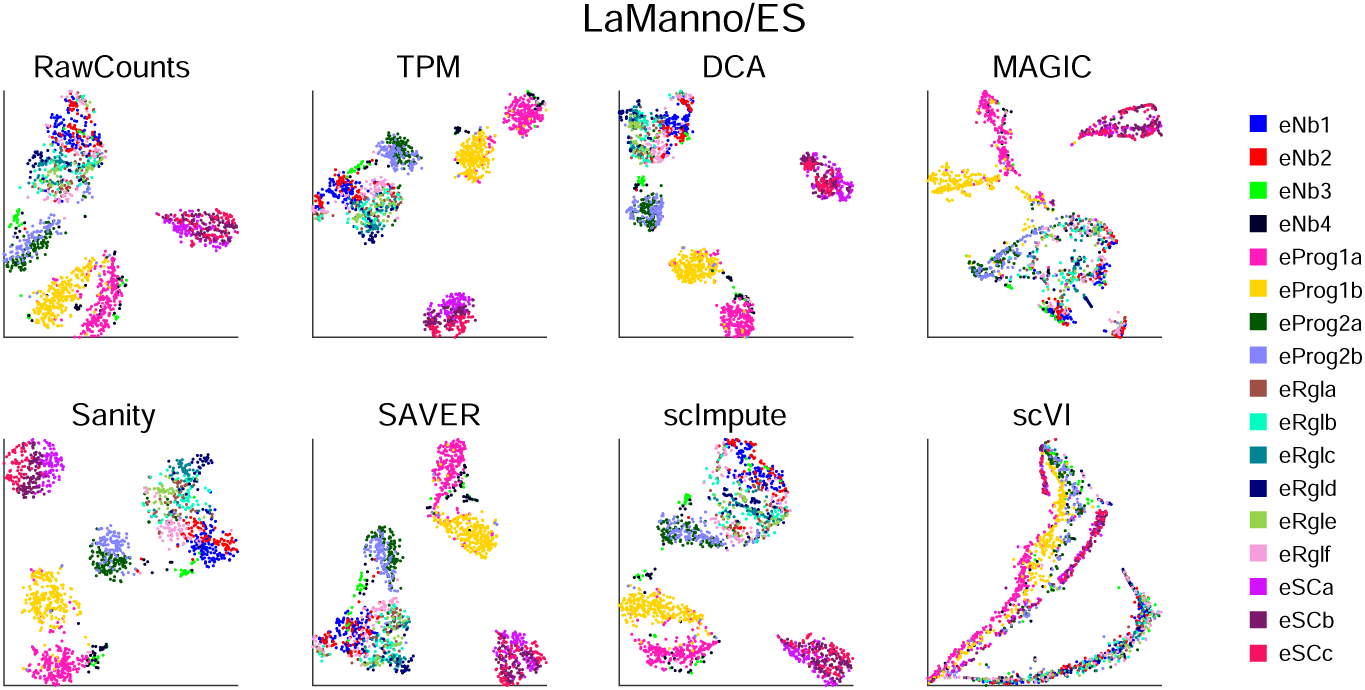
Each panel show a t-SNE visualization of the LaManno/ES dataset using the normalized gene expression values of the method indicated at the top. Each point represent a cell and is colored by the cell type

**Figure S9:**
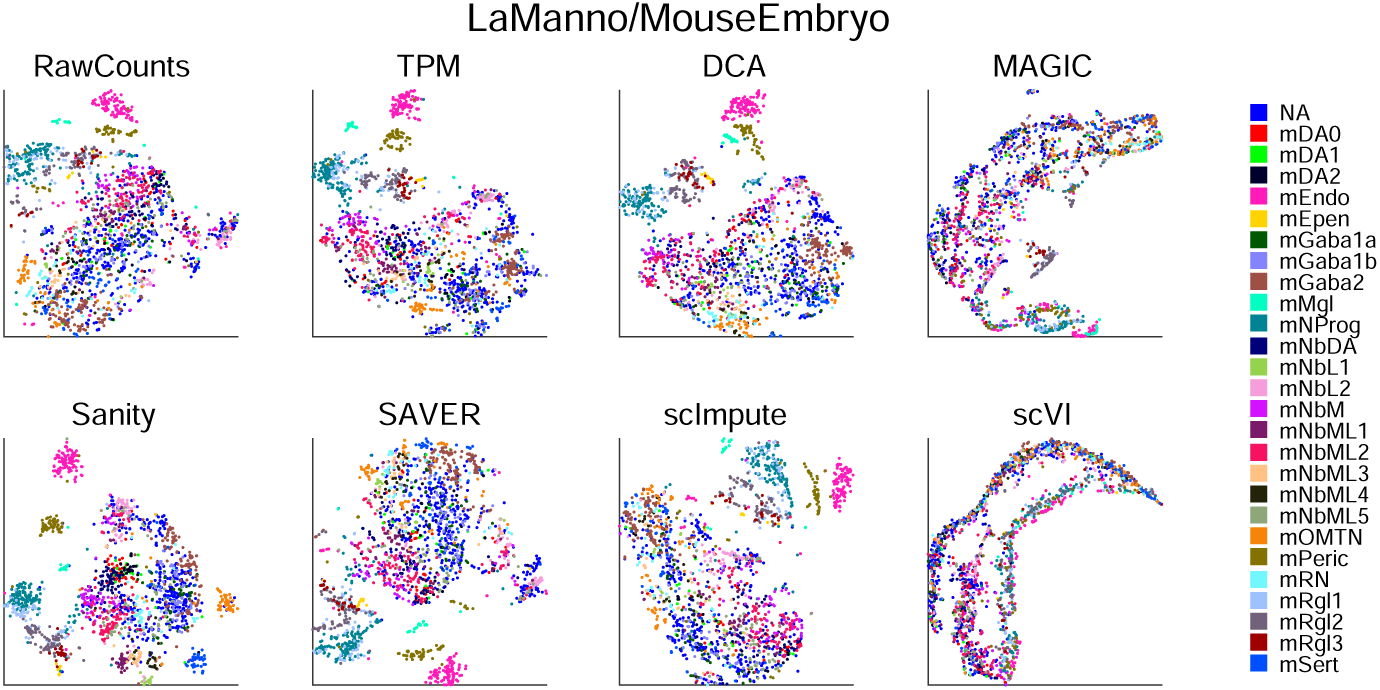
Each panel show a t-SNE visualization of the LaManno/MouseEmbryo dataset using the normalized gene expression values of the method indicated at the top. Each point represent a cell and is colored by the cell type annotated in the original publication.

**Figure S10:**
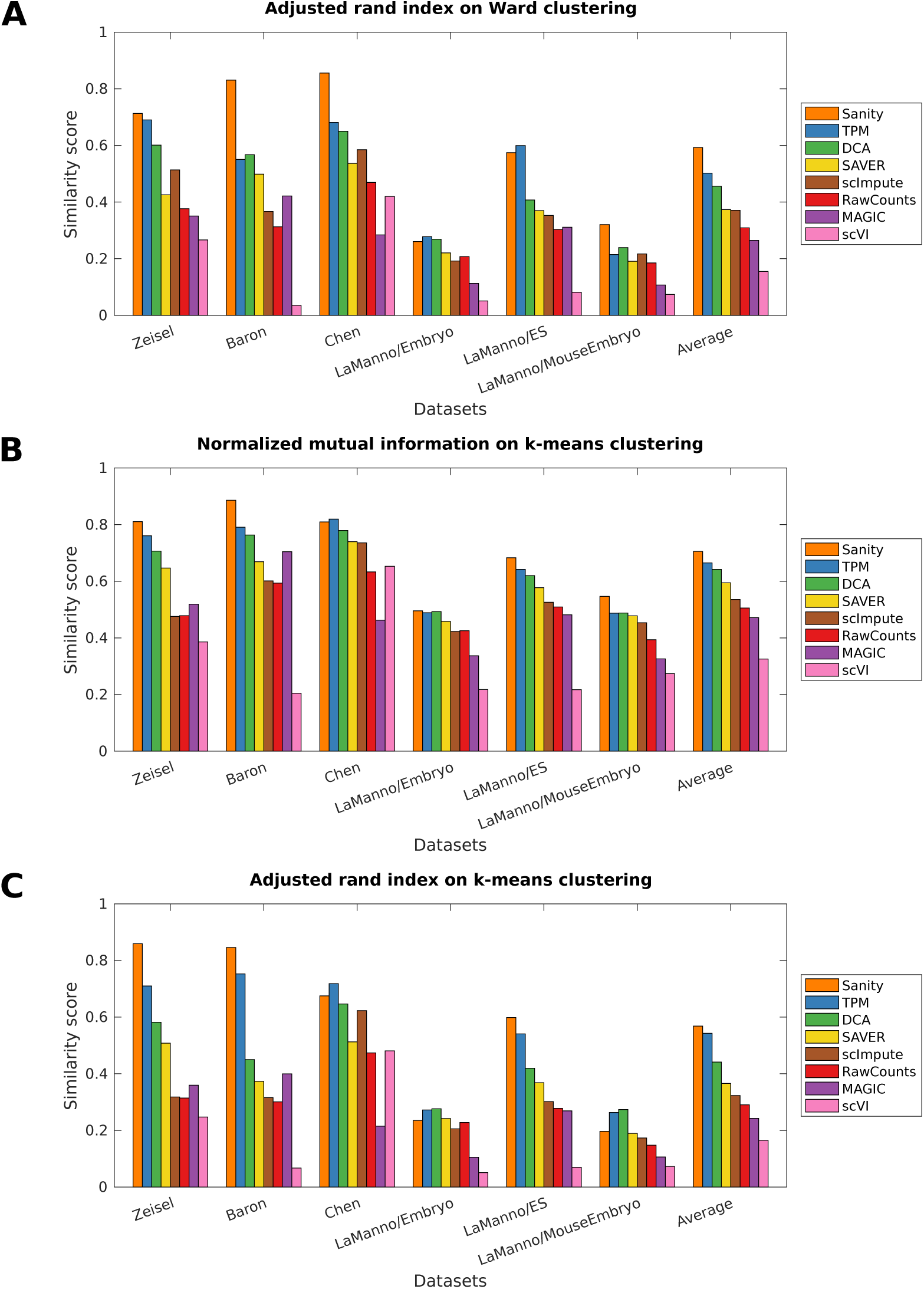
Similarity between the reference clusters and the clusters inferred using the normalized gene expression values of the different methods. Clustering was carried out using either hierarchical clustering with Ward’s method (**A**) or using k-means clustering (**B** and **C**). The similarity measures used were the Adjusted rand index (**A** and **C**) and the normalized mutual information (**B**). Both measures take values between 0 (no similarity) and 1 (perfect similarity). Each group of bars shows the results for a particular dataset indicated below) and colors indicate the different methods (see legend). The last group of bars shows the average similarity per method across all datasets. Methods are sorted from left to right according to their average similarity.

**Figure S11:**
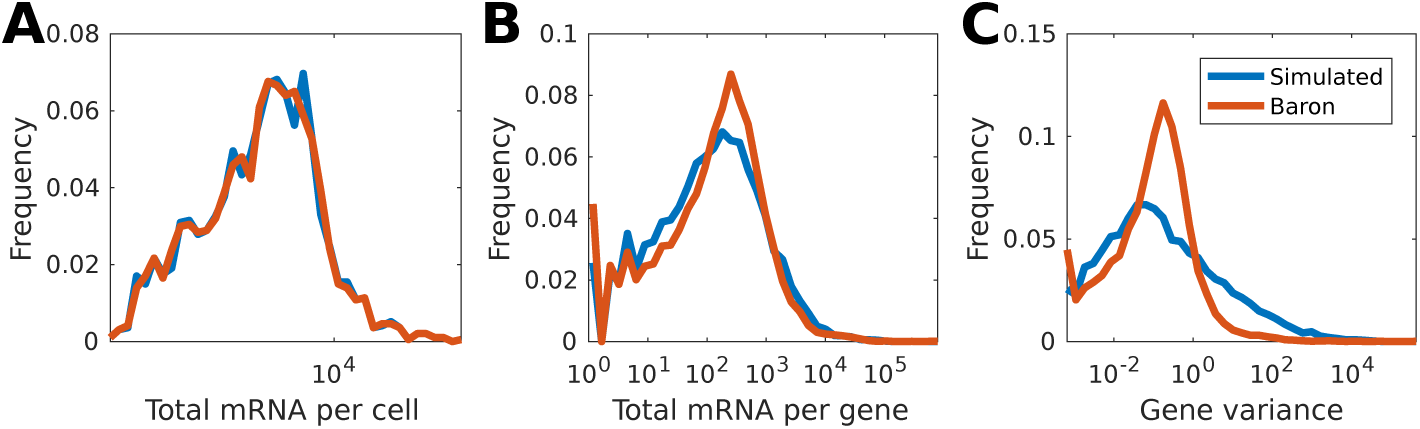
**A** Distribution of total mRNA captured per cell in the simulated dataset (blue) and the *Baron* dataset (red). **B** Distribution of total mRNA captured per gene in the simulated dataset (blue) and the *Baron* dataset (red). **C** Distribution of variance per gene calculated on the raw count matrix obtained from the simulated dataset (blue) and the *Baron* dataset (red).

